# Energetic costs of cellular and therapeutic control of stochastic mtDNA populations

**DOI:** 10.1101/145292

**Authors:** Hanne Hoitzing, Payam A. Gammage, Michal Minczuk, Iain G. Johnston, Nick S. Jones

## Abstract

Mitochondrial DNA (mtDNA) copy numbers fluctuate over time due to stochastic cellular dynamics. Understanding mtDNA dynamics and the accumulation of mutations is vital for understanding mitochondrial-related diseases. Here, we use stochastic modelling to derive general results for the impact of cellular control on mtDNA populations, the cost to the cell of different mtDNA states, and the optimisation of therapeutic control of mtDNA populations. We provide theoretical evidence that an increasing mtDNA variance can increase the energetic cost of maintaining a tissue, that intermediate levels of heteroplasmy can be more detrimental than homoplasmy even for a dysfunctional mutant, that heteroplasmy distribution (not mean alone) is crucial for the success of gene therapies, and that long-term rather than short intense gene therapies are more likely to beneficially impact mtDNA populations. New experiments validate our predictions on heteroplasmy dependence of therapeutic outcomes.

## 1 Introduction

Most human cells contain 100–10,000 copies of mitochondrial DNA (mtDNA) which are situated inside the mitochondria. The proteins encoded by mtDNA are crucial for mitochondrial functionality, and mutations in mtDNA can cause devastating diseases [1, 2, 3, 4, 5, 6]. Heteroplasmy, the proportion of mutant mtDNA molecules in a cell, has to pass a certain threshold (typically 60–95%) before any biochemical defects can be observed [7, 8, 9, 10, 11, 12, 13]. Evidence that mild mutations, that are not obviously pathological, can have physiological effects is also appearing [14]. The existence of thresholds at which mutant loads begin to have an effect has profound implications for our understanding of disease onset, drawing attention to the variance dynamics of the mutant fraction in cellular populations. As this variance increases more cells can be above threshold, and thus show pathology, *even if average mutant load is unchanged*.

How cellular population fractions of mutant mtDNA change over time is not well understood. The cell exerts control on mitochondrial populations in response to changes in mtDNA copy numbers [15, 16]. Mitochondrial biogenesis and maintenance require cellular resources, and mitochondria are key sources of ATP and play other important metabolic roles: a tradeoff of bioenergetic costs and benefits is thus involved in the interaction between the cell and its mitochondria. The particular ‘effective cost’ that cellular control acts to minimise remains poorly understood: for example, both decreases [17] and increases [17, 18] in wildtype copy numbers have been observed for different mutations as the mutant load increases. Some studies suggest that mtDNA density is controlled [19, 20, 21], others that total mtDNA mass [22, 23], or mtDNA transcription rate [24] is controlled. Understanding the dynamics of mtDNA populations inside cells, and how these populations react to clinical interventions, is crucial in understanding genetic diseases [25, 26]. However, experimental tracking of mtDNA populations over time is challenging, necessitating predictive mathematical modelling to provide a quantitative understanding of these systems.

In parallel with efforts to elucidate cell physiological control, protein engineering methods to control mtDNA heteroplasmy are making fast progress. Two recently developed methods for cleaving DNA at specific sites involve zinc finger nucleases (ZFNs) and Transcription Activator-like Effector Nucleases (TALENs) [27, 28, 29, 30, 31, 32, 33], which have been re-engineered to specifically cleave mutant mtDNA [34, 35, 36, 37, 38]. Mito-TALENs have been successfully used to reduce mutant loads in cells containing disease-related mutations, but elimination of the target mutant mtDNA was not complete [39, 34]. Similarly, treating cells multiple times with mtZFNs led to near-complete elimination of mutant mtDNAs [37, 38]. Quantitative theory for these promising therapeutic technologies has not yet been developed, leaving open questions about how these tools can be optimally deployed.

In this paper, we develop theory from bottom-up bioenergetic principles which allows us to study the effects of distinct cellular mtDNA control strategies (Section 2.1), to analyse the bioenergetic cost of different mtDNA states (Section 2.2), and to combine mtDNA control and energy-based cost (Section 2.3.1) to identify optimal control strategies for the cell. Finally, we construct a model for artificial mtDNA control using recent experimental data [38] to propose optimised treatment strategies while highlighting challenges linked to heteroplasmy variance (Section 2.3.2).

## 2 Results

### 2.1 Control: general insights on the role of feedback control

We employ the ‘relaxed replication’ model for heteroplasmic mtDNA populations (populations in which wildtype *(w)* and mutant (*m*) mtDNA molecules co-exist). Each mtDNA molecule replicates and degrades according to Poisson processes with rates λ and *µ* respectively [40, 41]. Because control of biogenesis or autophagy yield similar behaviours [42], we assume that the degradation rate µ is constant and that feedback control is manifest through replication rate (λ = *λ*(*w,m*)). To connect with experiments, we use *µ* ≈ 0.07 day^−1^ corresponding to a half-life of about 10 days [43]. We consider the case where no selective difference exists between mtDNA types, although our theory is readily generalised to include such differences.

#### A wide range of control strategies induces similar mtDNA behaviour and admits quantitative analysis

Whatever the quantity being controlled, in healthy cells the intuitive aim of homeostatic mtDNA control is to guarantee cell functionality by keeping the wild-type number around a particular value and within certain bounds. Many possible control strategies can be parameterised to give rise to a specific wildtype distribution; also when mutants are present the means and variances of wildtype, mutant and heteroplasmy can be nearly identical for these differing strategies up to long times (~ 80 years) (Figure 1A,B,C, Figure S1B,C,D). It is not the manner in which the controlled quantity is being controlled, but *which* quantity is controlled that is the most important (Figure 1D).

We stress the difference between two types of average heteroplasmy (as was also mentioned in [41]): the individual cellular mean heteroplasmy 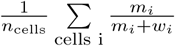 and the tissue homogenate heteroplasmy 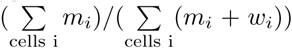. The difference between individual and homogenate means is clearly seen in Figure 1C. A tissue can thus appear, when studying the homogenate heteroplasmy, to show selection for one type of mtDNA over another, whereas in fact mean cellular heteroplasmy is unaltered and both mutant and wild types have the same proliferation rates.

From here on we focus on a linear feedback control of the form *λ*(*w,m*) = *c*_0_ – *c_1_(w* + *δm*), with *c*_0_ > 0, *c_1_* > 0 and *δ*, corresponding to the strength of sensing of mutant mtDNA, constants. The deterministic steady states of this system, (*w_ss_,m_ss_*) are given by 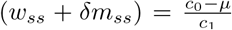, with *µ* the constant degradation rate. This line of constant *(w_ss_* + *δm_ss_*) forms a straight line in (*w, m*) space and stochastic dynamics will fluctuate around these steady state lines, with corresponding changes in heteroplasmy (Figure S1A). The parameter *c*_1_ determines the control strength and *δ* allows for distinct control strengths for wildtypes and mutants.

#### Variance behaviour over time in mtDNA populations

Van Kampen’s system size expansion can be used to predict mtDNA variance (Table 1E), showing good agreement with stochastic simulations up to hundreds of days [42]. Applying this approximation to a general form of mtDNA control, we here find that *i) if only one species is controlled, the variance of the controlled species reaches a constant value (see also [42]), ii) when both species are controlled with equal strength their variances increase at identical rates*, ii*i) in general the more tightly controlled species has a more slowly increasing variance, and iv) the rate of increase of hetero-plasmy variance depends, to first order, only on mtDNA copy number and turnover (as found in [42]).* (Figure 1, Table 1(I)).

**Figure 1:**
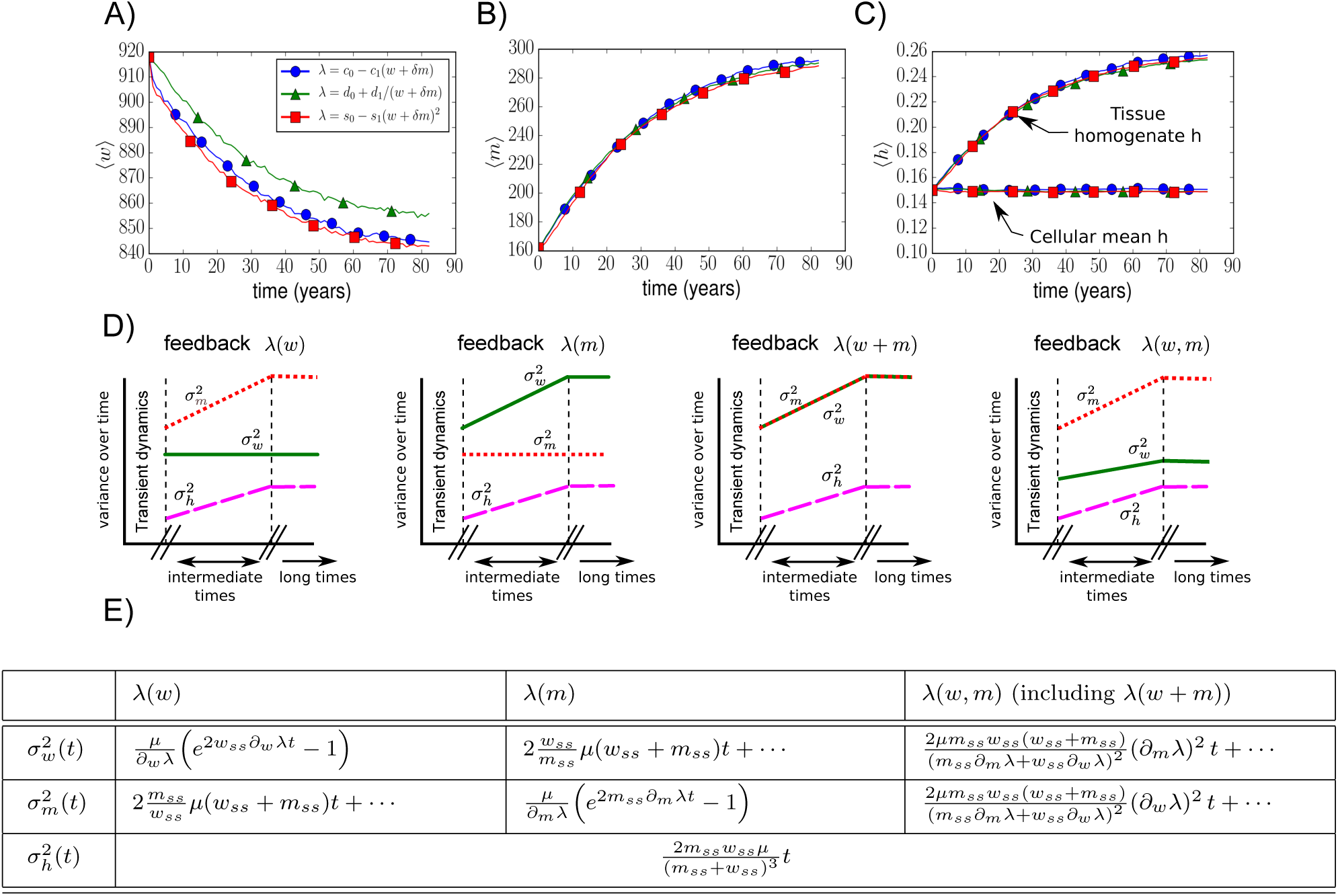
A) – C) A wide range of cellular control strategies can yield similar dynamics. Stochastic simulations were used to compare three structurally distinct cellular controls (see legend), each reflecting a different function of the underlying sensed quantity *w* + *δm* with *δ* = 0.5. All controls are set to have the same wildtype mean and variance in the absence of mutants (Section S2). No marked difference in mean and variance behaviours of wildtype, mutant, and heteroplasmy were seen in simulations up to ~ 80 years (see also Section S2). Figure (C) illustrates the difference between cellular mean heteroplasmy 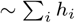, which remains constant, and tissue homogenate heteroplasmy 〈*m〉*/(*〈m〉* + (*〈w〉*), which increases over time (if *δ* > 1 it would decrease over time). **D) Control tradeoffs are required when multiple species are** present. The more strongly one species is controlled, the more control is lost over the other. Changes in variances as described by the Linear Noise Approximation (see Section S1) are shown (intermediate times). For long times, extinction of one of the species is likely and the variance of the surviving species saturates. For the control *λ*(*w,m*) we have depicted the case in which mutants contribute less to the control than wildtypes. **E) Analytical expressions for the means and variances according to the Linear Noise Approximation.** Solutions are shown for wildtype, mutant, and heteroplasmy variances for various types of control. Dots indicate constant or exponentially decaying terms; full solutions are given in Section S1. Note that the initial rate of increase of heteroplasmy variance does not depend on the control specifics, but only on mtDNA copy number and turnover (see also [42]).

**Table 1:** Here we present key results of the paper, which hold under the assumptions used in our models. We model mtDNA dynamics using stochastic birth-death simulations and assume: cells are heteroplasmic (containing both wildtype and mutant mtDNA molecules); birth and death rates are identical for wildtype and mutant species; the mtDNA dynamics are subject to cellular feedback control. We introduce a mitochondrial energy-based cost function; results referring to optimal controls and costs depend on the structure of this cost function which is discussed in Section 3 and Section S4.

|  |  |
| --- | --- |
|  | Key results presented in this paper. |
| I | If only one mtDNA species is controlled the variance of the controlled species reaches a constant value. When both species are controlled with equal strength their variances increase at identical rates, and, in general, the more tightly controlled species has a more slowly increasing variance (Figure 1, Section 2.1). |
| II | The mean energetic cost of maintaining a tissue can increase over time due to the nonlinear influence of mtDNA variance, even if the energetic demand on the tissue stays the same and mean levels of mtDNA are constant (Section 3, Eq. 3). |
| III | A control lacking any mutant contribution shows an exponentially increasing cost, and the effects of particular cellular control strategies are more pronounced in low copy number cells (Section 2.3.1, Figure 3A, B). |
| IV | Control strategies based on the energy status of the cell will often outperform control based on mtDNA copy number or sensing mtDNA mass (which would work well for deficient deletion mutants, but would be suboptimal for deficient point mutations) (Section 2.3.1, Figure 3C). |
| V | Even for pathological mutants, reduction of mutant mtDNA alone is not always the optimal control strategy for a cell to adopt (Section 2.3.1, Figure 4). |
| VI | Tissues with high mean heteroplasmy levels will generally be harder to treat with mitochondrially targeted endonucleases if the heteroplasmy variance is high, especially if this high mean level is caused by a small percentage of cells (Section 2.3.2, Figure 5D). |
| VII | Weak long-term rather than short intense endonuclease treatments are more likely to beneficially impact mtDNA populations (Section 2.3.2, Figure 5G, H). |

If a given mtDNA state (w, m) accrues a cost *C(w, m*) to the cell (this could be an energetic cost or some other metric of tissue burden), increasing variances in *w* and m can lead to the mean cost 〈C(w, m)〉 changing even when mean cellular copy numbers 〈w〉, 〈m〉 remain constant (Methods), as the mean of a nonlinear function of random variables is not generally equal to the function of the mean of those variables (as seen above with cellular vs homogenate heteroplasmy). *The mean cost of maintaining a tissue may thus increase over time, even if the tissue demands stay the same and mean levels of mtDNA are constant* (Table 1(II)). However, these increases may be small and whether they are significant depends on the details of the cost function: hence the need to consider the explicit forms in the next section.

### 2.2 Cost: an effective mitochondrial energy-based cost function

Here, we attempt to build a cost function that assigns a cost to a given mtDNA state (*w*, *m*) and allows a general quantitative investigation of the tradeoffs in maintaining cellular mtDNA populations. The ‘true’ energy budget of a cell with a given mitochondrial population is highly complex, involving many different metabolic processes in which mitochondria are involved [44, 45, 46]. We provide a simpler description, focussing on ATP production as a central mitochondrial function, and removing kinetic details in favour of a coarse-grained representation, to provide qualitative rather than quantitative results and provide descriptions of its connection to biological observables (Methods and Section S4).

#### General cost function structure

Three important terms involved in the energy status of a cell are: i) the energy demand *D*, ii) the net energy supply *S*, and iii) the efficiency with which the energy is supplied. Here we define efficiency as the amount of energy that is produced per unit of resource consumed. For this study, we focus on mitochondria as the central bioenergetic actors in the cell, with wildtype and mutant mitochondria consuming resource and producing energy currency.

We express our effective cost function as:

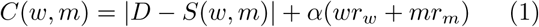

where α is a constant, and *r_i_* gives the rate of resource consumption of a mitochondrion of type *i (w* or *m*). The second term assigns a cost to the usage of resource. The terms in this cost function are expressed as rates: *S* and *D* then correspond to energy production (supply) and demand per unit time. This cost function assigns the lowest cost to a state that satisfies demand in the most efficient way, and both an excess or deficit of supply are penalized.

Energy production, *S(w, m*), is modelled as

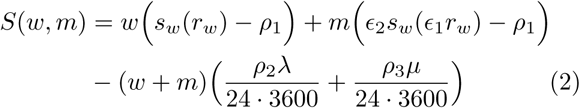

where ρ_1_,_2,3_ are mitochondrial maintenance, building, and degradation costs, *s_w_(r_w_)* denotes the energy production rate (per second) for a single wildtype mitochondrion given a resource consumption rate *r_w_* (Methods, Section S4), and λ and *µ* denote the birth and death rates in units day^−1^. Mutant mtDNA molecules are distinguished by the parameters ϵ_1_, *ϵ*_2_ ∈ [0,1] describing the mutant resource uptake rate (ϵ_1_) and efficiency (ϵ*_2_*) relative to that of the wildtypes. The lower the value of ϵ_1_, the lower the mutant energy output as a consequence of less resource consumption.

Despite the coarse-grained effective nature of our cost function, plausible parameterisations can be estimated (Table S1); further details on the choice of parameter values, and their biochemical interpretations are given in Section S4. We consider two different types of cost function (Methods, Section S4): the relationship between the mitochondrial resource consumption rate and energy production rate is assumed to be i) linear, or ii) saturating. The different systems will be referred to as ‘the linear output model’ or ‘the saturating output model’. The saturating output model assumes that mitochondria eventually become less efficient as the flow through their respiratory chain increases. For both these models we use two different cost function parameterisations: one for high and one for low copy number cells.

One particular parameter we will use throughout this paper is *w_opt_*, the optimal (i.e. cheapest) number of wild-type mtDNA molecules in the absence of mutants. Each of our four different systems (saturating and linear, for low or high copy numbers) has its own value for *w_opt_*(Table S1).

#### Intermediate heteroplasmies may be inefficient and resource availability can dictate the cost of mtDNA states

Figure 2 shows the value of the cost function in *(w, m*) space, for different mutant pathologies (different values of ϵ*_1_*). Interestingly, it is possible to satisfy demand with only wildtype or only mutant mtDNAs, but not at certain intermediate heteroplasmies. This counter-intuitive result aligns with arguments that mixed mtDNA populations are disfavoured [14] and can be energetically explained as below (referring to Figure 2B).

Increasing heteroplasmy while keeping total copy number constant can be interpreted as replacing a wildtype mitochondrion with a mutant, leading to a tradeoff. The changed mitochondrion (from wildtype to mutant) produces less energy than before; all other mitochondria need to consume more resource to maintain a constant total output, and so become less efficient due to output saturation. However, the changed mitochondrion itself has become more efficient because of its lower resource consumption rate. The tradeoff between these two factors depends on *h* itself; a more detailed discussion is given in Section S5. *The result is that intermediate heteroplasmies are the least efficient and the most expensive* (Figure 2 and S3). We note that this is only the case if i) at high respiration rates, energy production becomes less efficient, and ii) mutants consume less resource than wildtypes (perhaps through ETC deficiency), effectively making them more efficient than wildtypes (due to output saturation). A model in which mitochondrial output always increases linearly with resource consumption does not show this behaviour (Figure 2C).

### 2.3 Combining cost and control: comparison and optimisation of both cellular control and treatment strategies

#### 2.3.1 Timescales and energy sensing in optimal control of mtDNA populations

Here we compare the mean cost over time of four different plausible cellular control strategies. The first two consist of the linear feedback control discussed earlier (λ(*w, m*) = *c*_0_ – *c_1_(w* + *δm))* with i) *δ =* 0 (only wild-types are sensed) and ii) *δ =* 1 (total mtDNA copy number is controlled without differentiating between mutant and wildtype). We also consider control strategies for the cell that are optimised under our imposed cost function. We identify the optimal parameterisations that minimise steady-state cost for (iii) a linear feedback control and (iv) the relaxed replication model [40, 41].

To optimise a control, both an optimization time-scale *T* and a set of initial conditions are required. Here we use *T = ∞*, corresponding to the steady state limit, and a set of initial conditions with heteroplasmies in the range *h_o_* ∈ [0,0.2]; we later consider finite values of *T.* We assume that in the absence of mutant mtDNA, each strategy is optimised and all have equal wildtype means and variances (Section S6); we compare the changes in dynamics induced by the presence of mutants.

Figure 3 shows the mean cost up to ~ 80 years resulting from stochastic simulations; cost variance, as well as mutant and wildtype dynamics were also computed (Figure S4). The relaxed replication rate control and the linear control λ(*w, m*) = *c_0_ – c_1_(w* + *δm)* behave very similarly when *δ* and γ take their optimal values. A control with *δ =* 0 shows an exponential increase in cost, though it still takes ~ 12 years and ~ 65 years for this control to become 10% more expensive than the others in our low and high copy number cells, respectively. We conclude that: *i) a control lacking any mutant contribution only becomes notably costly on the order of 10 years, and ii) effects of particular control strategies are more pronounced in low copy number cells* (Table 1(III)).

We now investigate how the optimal value of mutant sensing *δ, δ_opt_*, for the linear control λ(*w, m) = c_0_ – c_1_(w* + *δm)* depends on timescale *T*, initial heteroplasmy *h_0_* and the ‘mutant pathology level’ described by parameter *ϵ_1_*. Biologically, this question reflects how the cell should optimise its processing of mtDNA state as the mutant load and severity changes. Intuitively, values of *ϵ_1_* ≃1 have *δ_opt_* ≈ 1: when wildtypes and mutants are equivalent, having a steady state with *w* + *m = w_opt_* is desirable.

Values for *δ_opt_* were found for the linear and saturating model, with low and high initial heteroplasmy values, for *T =* 100 days (Figure 3C). Having *δ* ≈ 1 means wild-types and mutants are fed back similarly, whereas δ ≪C 1 means mutants are fed back much less. For very deficient mutants (low *ϵ_1_*), a low *δ_opt_* ensures that wildtype copy number remains close to its optimal value to compensate for the mutants. Generally, as *ϵ_1_* decreases, *δ_opt_*decreases. In the linear model *δ_opt_* becomes negative for low *ϵ_1_* values; as mutant copy number increases, a negative *δ* leads to an increase in wildtype to compensate for the deficient mutants. Similar results are found for longer timescales *T* (Section S6).

When mitochondrial energy outputs are sensed, a deficient mutant will contribute less, so a low (or high) value for *ϵ_1_*is automatically associated with a low (or high) value of *δ.* We can see that *control strategies based on the energy status of the cell will often outperform control based on mtDNA copy number (which always has δ = 1) or sensing mtDNA mass (which would work well for deficient deletion mutants, but would be suboptimal for deficient point mutations)* (Table 1(IV)).

##### Locally optimal control strategies map the control space of mtDNA populations

With the use of our cost function it is possible to identify *locally optimal controls:* controls that, for each state (*w, m*), move the system in the direction of the largest decrease in cost (Figure 4).

When heteroplasmy is high the main priority is not always to decrease mutant copy number, but to increase wildtype copy number even if this means an increase in mutant load (region 2 in Figure 4A). It is only after wildtype copy number has sufficiently increased that the focus should be on decreasing *m.* At high copy numbers the optimal dynamics are to decrease all mtDNA in an evenhanded manner (region 1) rather than decreasing m at a faster rate than *w.* For the saturating model, we also observe a divergence point in the space of local optimal strategies, reflecting the two local cost minima (high wildtype and high mutant) observed earlier (Figure 2). Hence, there are several regions of state space where *even for pathological mutants, reduction of mutant mtDNA alone is not always the optimal control strategy* (Table 1(V)). Finally, the less pathological the mutants become (e.g. Figure 4B), the more the locally optimal control starts to resemble a linear control. In the linear output model, the optimal control is always quite linear (Figure 4C).

**Figure 2:**
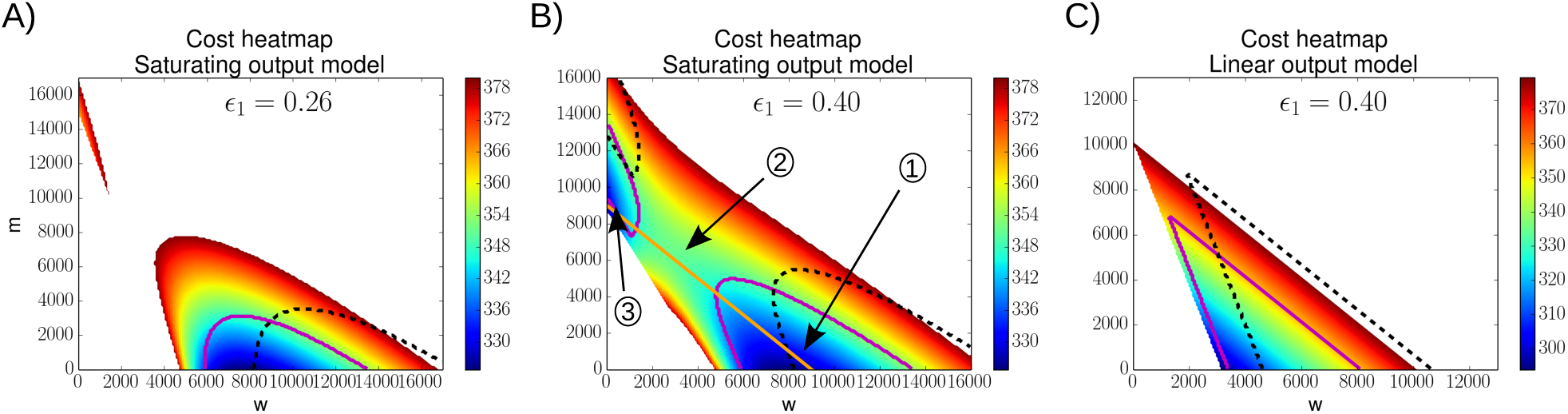
Intermediate heteroplasmies can be less efficient than either wildtype or mutant homoplasmy. A visualization of the cost function in (*w,m*) space is shown for the saturating and linear mitochondrial output model. Only the cost inside the demand-satisfying region is shown, for various mutant pathologies; hence, white regions correspond to states where cellular demand is not satisfied though cells may survive by, for example, increasing glycolysis, effectively reducing mitochondrial demand. All the plots in this figure are at high copy numbers, results are qualitatively similar for low copy numbers. **A)** The magenta (solid) and black (dashed) lines show the contour of the demand-satisfying region when demand is increased by 10%, or demand is increased by 50% and cellular resource availability is increased by 35%, respectively. Fluctuations increasing demand by 10% can significantly reduce the demand-satisfying region; larger fluctuations can only be handled when total cellular resource uptake rates increase. **B)** The orange line corresponds to constant total copy number; moving up along this line increases heteroplasmy. Cells in region 1 or region 3 are more efficient, and show a lower cost, than cells in region 2. **C)** The linear mitochondrial output model does not show a decreased efficiency at intermediate heteroplasmy values.

**Figure 3:**
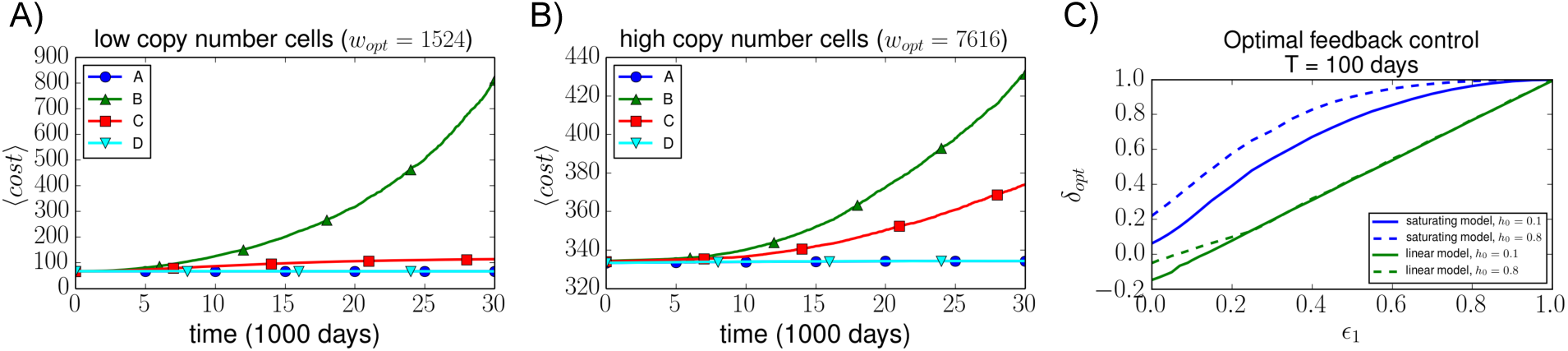
A control that senses no mutations shows an exponentially increasing cost, which is most noticeable in low copy number cells. Here we show the mean cost for the following four controls: A) the optimised ‘relaxed replication control’ 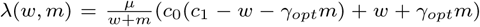 [40], and linear feedback controls λ(*w*, *m)* = *c*_0_ – *c_1_ (w* + *δm)* with B) δ = 0, C) *δ=1*, and D) *δ* = δ_o_*_pt_.* The controls were initialized in steady state at *ho* = 0.15 and simulated for ~ 82 years (30000 repeats). Both figures used the saturating output model; the left and right figures correspond to low and high copy number cells respectively. The free parameters left in control A and D were optimised over initial conditions in the range *h* ∈ [0,0.2]. Here ϵ_1_ = 0.3 was used, in the section below we investigate more values of ϵ_1_. Other control parameters used are given in the SI. **C) A control based on sensing mitochondrial energy output may be generally a good strategy**. This plot shows the optimal value of *δ* in the control *λ*(*w,m*) = *c*_0_ – *c_1_(w* + *δm)* as a function of *ϵ_1_*, for the linear and saturating model and for both low (*h_0_* = 0.1, solid line) and high (*h_0_* = 0.8, dashed line) initial heteroplasmies. Here we used *T* = 100 and high copy number values for both models. The optimal mutant control contribution is broadly lower when mutants are more deficient. Similar plots for *T* = 10^4^ are shown in the SI.

**Figure 4:**
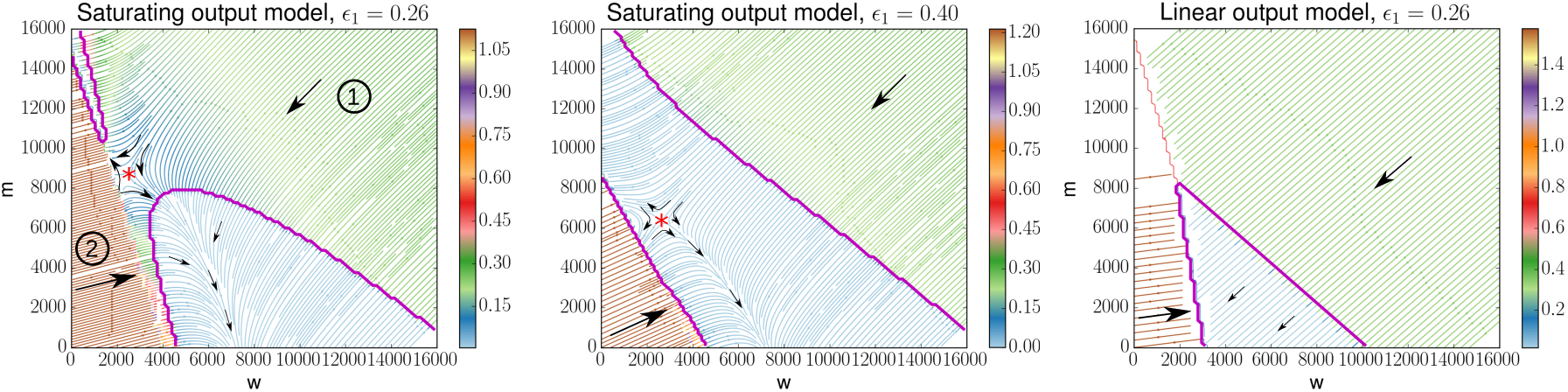
Locally optimal controls show non-linear behaviours close to the demand-satisfying regions, but linear optimal dynamics far away from these regions. Here we have used streamplots to visualize the locally optimal control in *(w, m)* space, for various parameters of ϵ_1_. At each point in the (*w,m*) plane, arrows show the direction of the optimal change to make to decrease cost *C(w,m).* Regions are coloured according to the magnitude of the decrease in cost when moving in the optimal direction. Black arrows illustrate general trends in these regions. A) Region (1) shows that at high copy numbers, both mutant and wildtype mtDNAs should be decreased in an evenhanded manner; region (2) shows the possibility that the optimal control involves an increase in mutant copy number. Here the saturating output model is used. B) Like figure (A), the saturating output model is used; a higher value for the parameter ϵ_1_ is used, meaning mutants are less pathological. C) Here the linear output model is used; the locally optimal control more closely resembles a linear control. In both (A) and (B) we see a divergence point (red asterisk) illustrating the fact that both high mutant and high wildtype states constitute local attractors of low cost (as in Figure 2).

### 2.3.2 A parameterised model of artificial mtDNA control for disease treatment

While the locally optimal controls identified above may not be achievable by the cell (for example, the cell may not be able to decouple biogenesis of wildtype and mutant mtDNA), genetic technology gives us the ability to artificially exploit these optimal strategies. Mitochondrially targeted Zinc Finger Nucleases (mtZFNs) [47, 48, 49, 37, 38] and mitoTALENs [34, 36] are able to produce shifts in heteroplasmy by specifically cutting mutant mtDNA; these technologies thus offer the prospect of gene therapeutic treatments for some mitochondrial diseases. Though results are promising, treatments with endonucleases are not perfect and can have substantial dose-dependent off-target effects [38]. To develop predictive quantitative theory to understand and tune the effects of these interventions, we model nuclease transfection as inducing selective increases in mtDNA degradation, on the background of the cellular feedback control described above (Methods). A fit (see Methods) of our model parameters describing strength (*Io*), duration (b), and selectivity (ξ) of nuclease treatment is shown in Figure 5A, B, illustrating the ability of this simple model to capture the complex dynamics resulting from nuclease activity. For every mutant that gets cut by the endonucleases, ξ wildtypes get cut (i.e. when ξ = 1 there is no distinction between mutants and wildtypes, when ξ = 0 there is no off-target cleavage). The best-fit value for the selectivity parameter ξ ≈ 0.78 implies a low nuclease selectivity; however, this may be due to the high mtZFN concentration that was used during the experiments (Discussion).

Nuclease treatment and a subsequent ‘recovery phase’ of cellular relaxation will have the net effect of mapping an initial heteroplasmy value *h_i_* to a mean final heteroplasmy value, *h_f_.* We simulated this mapping of initial to final heteroplasmy values in the presence of cellular feedback control (Figure 5), and observed that the *heteroplasmy shifts are similar in low and high mtDNA copy number cells (the variance of the shift is slightly lower for high copy number cells), and that the shift is largest for intermediate heteroplasmies.* Interestingly, *for high h values, it is possible to end up with a* higher *heteroplasmy value after treatment, especially if* ξ ≃ 1 (Figure S6).

#### Knowledge of the heteroplasmy distribution of a tissue is important in determinining how eff-ciently the tissue can be treated

To explore the effect of the heteroplasmy distribution on treatment efficacy, we consider three different initial *h* distributions with different heteroplasmy variances, but identical homogenate means. We treat these populations multiple times using the parameters fitted in the previous section, and observe the shift in heteroplasmy distribution as well as the change in heteroplasmy mean and the probability of crossing a pathogenic heteroplasmy threshold *P(h* > 0.6) (Figure 5D, E).

High heteroplasmy variances require many cells close to the two extremes *h =* 0 and *h =* 1, which are challenging to shift. A striking reduction in treatment efficacy is predicted as heteroplasmy variance increases while mean heteroplasmy stays constant (Figure 5D, E). Threshold crossing probability (for example, *P(h* > 0.6)) also becomes harder to shift at higher heteroplasmy variance. These differences in treatment efficacy depend on the value of *δ* (mutant sensing) in our model: lower *δ* will decrease the difference in efficacies. We conclude that *tissues with a high mean heteroplasmy levels will generally be harder to treat if the heteroplasmy variance is high, especially if this high mean level is caused by a small percentage of cells*, with the strength of this effect dependent on mutant sensing (Table 1(VI)). We tested our finding that it is difficult to shift high heteroplasmy values by looking at new data from experiments focussing on the treatment of high heteroplasmy cells and find that the new results support our theory: mtZFNs were transfected into 143B cells bearing 99% m.8993 T>G mtDNA, but no shift in heteroplasmy was measured 14 days post-transfection, nor at 28 days when mtDNA copy number had recovered to control level (Figure S7).

**Figure 5:**
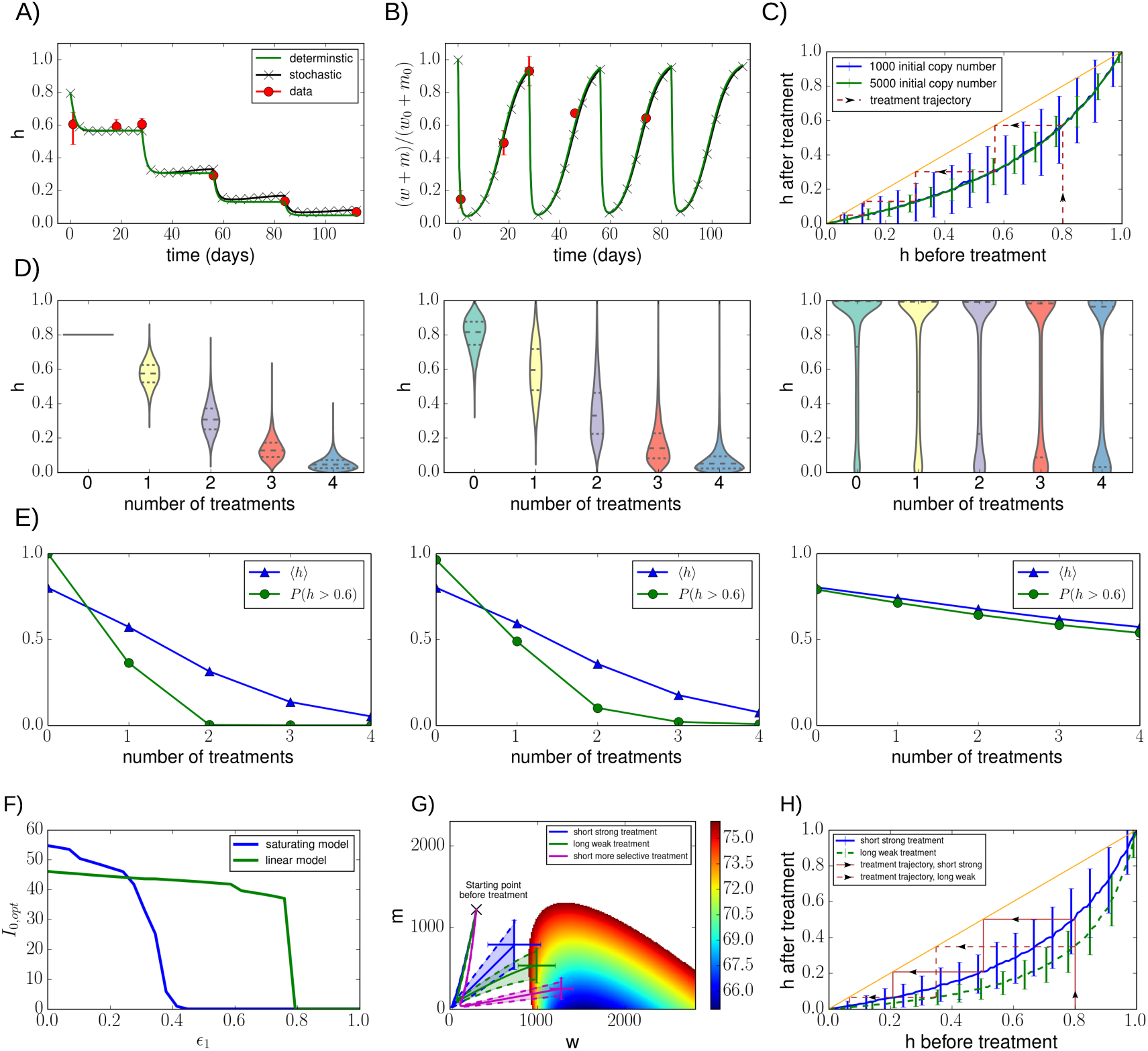
A simple model of treating cells with mitochondrially targeted endonucleases captures experimental observations of cellular mtDNA statistics. A, B) Our treatment model was used to fit recent experimental data [38], good agreement was found for both deterministic and stochastic simulations; details about the fitting produce are given in Methods. C) A given initial mean heteroplasmy value maps to a final mean value after a round of treatment and recovery. Stochastic simulations were performed for both low (blue, 1000) and high (green, 5000) copy number cells. The orange line denotes identity, and the purple trajectory shows the heteroplasmy shifts for multiple treatments starting at *h* = 0.8. Further details of the fitting procedure are given in Methods and Section S7. **Knowledge of the heteroplasmy distribution is important in predicting how efficiently a tissue can be treated. D, E)** The effect of four consecutive treatments on three different initial heteroplasmy distributions is shown; all initial distributions have identical means (〈*h*〉 = 0.8) but different variances (increasing from left to right). The higher the variance of the initial population, the harder to shift mean heteroplasmy values; mean values after each treatment as well as *P(h* > 0.6) are shown more clearly in figure (E). In these simulations we assumed that every cell gets transfected. **Gentle but sustained treatments induce larger heteroplasmy shifts than hard and brief treatments. F)** Both the linear and saturating model show a sharp drop in the optimal treatment strength *Io,_o_pt* as the mutants become more functional (i.e. as ϵ_1_ increases). G) Means and variances of mutant and wildtype copy numbers were simulated during a round of treatment and recovery, using: i) fitted parameters (blue), ii) a longer treatment duration (smaller b, green) and iii) a higher selectivity (smaller ξ, magenta). The longer weaker treatment induces higher heteroplasmy shifts than the shorter stronger treatment. Not surprisingly, a higher selectivity also leads to an improved heteroplasmy shift. Note that the variance of the final values is lower for more selective treatments. Error bars show standard deviations (based on 10 stochastic simulations), further detailed are given in Methods and Section S7. H) This figure again illustrates that gentle sustained treatments lead to larger heteroplasmy shifts. Examples of treatment trajectories are shown; after a single treatment, an initial heteroplasmy of 0.8 is mapped to 0.53 (short strong treatment) or 0.39 (long weak treatment).

#### Optimal clinical interventions

We can use our parameterised theory to find optimal treatment strengths *I_0,opt_* for a given system. Figure 5F shows *I_0,opt_* as a function of *ϵ_1_*. Intuitively, the strongest treatment should be given to the least functional mutants, and when mutants are almost as functional as wildtypes it is preferable not to treat at all. The optimal treatment strength drops rather sharply as *ϵ_1_*increases, and does so sooner for the saturating model. This last observation may be because at some point reducing heteroplasmy becomes more expensive as can be seen in Figure 2B. Optimal treatment strengths for longer treatments (higher *b)* show similar qualitative behaviour.

Figure 5G shows, using the identified values for *I_0,opt_*, the trajectories in (*w, m*) space throughout a single treatment and recovery phase; we used *ϵ_1_* = 0.3 and the corresponding cost heatmap is also shown. The three trajectories shown correspond to: i) a short and strong treatment (using the fitted parameter values), ii) a long and weak treatment, and iii) a short but more selective treatment. It can be seen that *treating longer but weaker results in a lower final heteroplasmy value than treating short and strong.* A weaker treatment also reduces the chance of a cell losing all its mtDNA molecules. A more selective treatment also leads to larger heteroplasmy shifts. The difference in treatment results for long compared to short treatments is also illustrated in Figure 5H. We note that in finding *I_0,opt_*, we initialized all cells in the same steady state. When a distributions of initial states is used, the variance that is now present is likely to affect the optimal treatment strength.

## 3 Methods

### Expected cost per unit time

Let the cost per unit time of state (*w, m*) be denoted by Λ, and the cost corresponding to the steady state *(w_ss_, m_ss_)* by Λ. Even if steady state copy numbers are constant over time (i.e. the mean values of *w* and m are always equal to *w_ss_* and *m_ss_)* the mean cost per unit time is generally not equal to Λ. By performing a Taylor expansion, the mean cost per unit time can be written as follows:

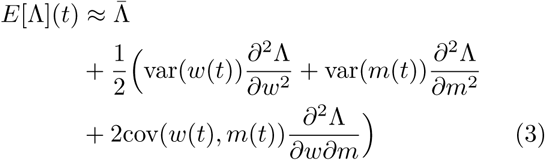

where E[Λ](t) is the expected cost per unit time given that the trajectory starts in state *(w_ss_*, m_ss_), and all partial derivatives are evaluated at steady state.

### Cost function structure

We assume that the net energy supply per unit time in a state *(w, m*), called *S(w, m*), involves the following four terms: (i) the *energy output* per unit time (*s_i_*) produced by the mitochondria; (ii) a *maintenance cost* per unit time (ρ*_i_)* to maintain the mitochondria, as their presence imposes some energetic cost (e.g. mRNA and protein synthesis); (iii) a *building cost (ρ_2_)* for the biogenesis of new mitochondria; and (iv) a *degradation cost (*ρ*_s_)* to degrade mitochondria. We will assume that every mtDNA molecule is associated to a particular amount of mitochondrial volume which we refer to as a ‘mitochondrion’ (Section S4).

At any time, mitochondria experience a certain energy demand and to meet this demand they need to have a certain resource consumption rate *r_i_* (where *i = w,m* refers to wildtype or mutant). Here we use the term ‘resource’ as an amalgamation of the substrates used for the oxidation system. We need to specify the relationship between the power supply (*s_i_*) and the rate of resources consumed (*r_i_*) by mitochondria. We use two different models *s_i_(r_i_*) which are justified in the SI:

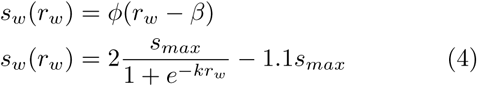

where *ϕ*, *β, k* and *s_max_* are constants respectively describing the mitochondrial efficiency, a basal proton leak-like term, the saturation rate of the efficiency, and the maximum energy production rate (Section S4).

We assume that pathological mutants can have a deficient electron transport chain (which may support a smaller flux leading to a lower resource consumption rate for mutants and therefore a lower ATP production rate) and a lower energy production efficiency:

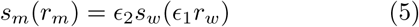

Here, *ϵ_1_*,*ϵ_2_* ∈ [0,1] describe the mutant resource uptake rate and the mutant energy production efficiency relative to that of a wildtype, respectively. In the main text we set *ϵ_2_* = 1; other values of *ϵ_2_* are discussed in Section S4.7.

The mitochondrial maintenance cost is denoted by ρ_1_ and corresponds to the energetic cost required to maintain the mitochondrion that contains the mtDNA. This energetic costs involves factors like the synthesis and degradation of mitochondrial proteins and enzymes. We assume the maintenance cost is the same for wildtype and mutant mitochondria (though for some mutations this is quite possibly not the case). The net energy supply per unit time, *S(w, m*), then follows as Equation 2.

To determine the value of *r_w_* for a given state *(w, m*), we first check whether the demand *D* (which we assume is a constant) can be satisfied in this state. If it can, we set equation (2) equal to *D* and solve for *r_w_*, i.e. we assume that if possible, the mitochondria will exactly satisfy demand. It may, however, not be possible to satisfy demand, which can be because of two reasons: i) there are not enough mitochondria present to produce enough energy, or ii) there is not enough resource available to meet demand. In the former case, we set *r_w_ = r_max_;* the mitochondria work as hard as possible to keep their energy output closest to demand. In the latter case, we assume that the total available amount of resource, *R* (which we consider to be constant), is shared equally between the mitochondria: 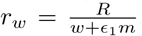. Cellular energy demand will naturally fluctuate over time, and because of excess capacity of mitochondria [50], there is no need to immediately increase mitochondrial biogenesis. In our model, a given state *(w, m*) may be able to satisfy a range of demands by changing its resource uptake rate. Further details of the cost function are given in the SI.

The parameters used in our cost function are summarized in Table S1, a discussion of the parameter values is also provided in the SI. Despite our model being simple, most parameters are biologically interpretable.

### Modelling control through mitochondrially targeted endonucleases

Experimentally, cells are transfected with two mtZFN monomers: one which binds selectively to mutant mtDNAs, and one that binds mutants and wildtypes with equal strength [49]. We simplify this picture by assuming an ‘effective’ mtZFN pool and use *[ZFN]* to denote its concentration. The increase in mtDNA degradation rate is then assumed to be proportional to *[ZFN].*

Nucleases are imported into the cell and then degrade over time, meaning that their concentration in the cell (and in the mitochondria) may be approximated by an Immigration-Death model. In the recent experiments [38], nucleases are expressed for short times which means that the immigration rate should be time-dependent and decreasing. This leads us to the following equation:

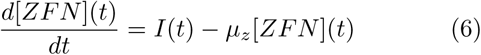

where *I(t)* and µ_Z_ are the immigration and death rates of the effective mtZFN pool, respectively. The immigration rate will increase sharply at the start of the treatment after which it decreases over time, and we have chosen to model *I(t)* as an exponentially decaying function, *I(t) = I_0_e^−bt^*, where *I_0_* is the initial rate right after the treatment is applied and *b* is a constant describing the duration of the treatment. The mtZFN concentration now becomes

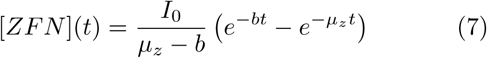

which is shown for various parameter values in Figure S6A. The mtDNA death rates at a time *t* after a treatment are then given by

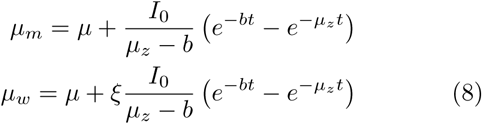

where *µ* denotes the normal baseline mtDNA degradation rate, 0 < ξ < 1 indicates how selective the treatment is (if ξ = 1 no distinction is made between *w* and m, if ξ = 0 no extra wildtype cleavage occurs during treatment).

To fit our model to recently obtained experimental data [38] we need to include the cellular feedback control which makes sure that copy number returns to its original value after the treatment. We will use λ(*w, m*) = *c*_0_ – *c_1_(w* + *δ m)* to model the feedback, and we assume δ *=* 1 because no significant changes in total mtDNA copy numbers were seen after treatments. We fitted the parameters *I_0_, b, ξ* and *c_1_.* Because mtDNA copy number in the cells used in the experiment is not known, we use both low (1000) and high (5000) initial copy numbers and fit our parameters for both cases (the value of *c*_0_ is set such that steady state copy numbers are 1000 or 5000). The obtained parameter fits are: *(I_0_, b*, ξ, *c_1_)* ≈ (38, 12, 0.76, 3 x 10^−4^) and *(I_0_, b, ξ, c_1_) ≈* (40, 12, 0.76, 5 × 10^−5^) for cells with initial copy numbers 1000 and 5000, respectively. For the mtZFN degradation rate we used a half-life of 1 day, to roughly match the experimental observation that almost no mtZFN was present 4 days post-transfection (with a half-life of 1 day, the size of an exponentially decaying population will be about 6% of the initial size).

### Shifting heteroplasmy in 143B cells bearing 99% m.899 T>G mtDNA

All methods used are as described in [37], more details are provided in Section S7.

## 4 Discussion

In this work, we have built a quantitative theory bridging stochastic optimal control, costs of mtDNA populations, and gene therapies. Our results contribute to a growing body of evidence [51, 52, 53, 54] that the variance of mtDNA populations has important physiological and therapeutic implications independently of mean heteroplasmy, and underline that stochastic theory is required to understand this biologically and medically important quantity. We compared different cellular strategies of regulating the number of mtDNA molecules, by studying the means and variance of the mutant and wildtype populations. We found that there exists a trade-off between controlling one species, or controlling the other. Heterogeneity in sensing may thus account for the large variety of mutant dynamics seen between different mutations and across different tissues. We introduced a mitochondrial energy-based cost function enabling us to compare distinct control mechanisms and found that a control based on the energetic status of the cell forms a good strategy in many situations. Recent experimental data [38] was used to fit a therapeutic treatment model in which cells are transfected with mitochondrially targeted endonucleases, with new experiments validating this theory. We showed that long gentle treatments induce higher heteroplasmy shifts than brief strong treatments, and that tissues with high heteroplasmy variances can be particularly hard to treat.

If the parameter *δ* is low, i.e. mutants are sensed less, mutant copy numbers at high heteroplasmies will be higher than wildtype copy numbers at low heteroplasmies. Experimentally it has been observed that heteroplasmic cells can have total mtDNA copy number values that are 5–17-fold higher than cells homoplasmic in wild-type [55, 56, 57, 58]. The cell has somehow allowed these mutants to expand, which may mean that they are less tightly controlled; Controls based on total energy output or mtDNA mass (which can result in *δ* < 1) may lead to such behaviours. A control on mtDNA mass could explain why deletion mutants are often seen to expand [59, 60] and would also predict normal copy number levels in cells harbouring mtDNA point mutations. Recently, it was found that samples with mtDNA indels had very high mtDNA copy number levels, but single nucleotide variants did not [61]. The cellular control mechanism could also be a mixture of copy number control, mtDNA mass control and energy sensing, meaning that mutations with different functionalities and masses all have different values for *δ* and expand to different extents.

We showed that heteroplasmy distributions in cell populations can provide important information about the possibility of successfully treating these cells. A tissue may be harder to treat if its high mean heteroplasmy level is caused by a small percentage of dysfunctional cells; these cells may also be hard to target using gene therapy if the transduction efficiency is low. Experimental values of mean homogenate heteroplasmy in heart tissue of patients with the 3243A>G mutation are roughly around 0.8, though ranges can be large [62, 63, 64, 65]. Muscle tissue often shows mosaic structures, with deficient patches of cells adjacent to healthy cells. These examples show that it may be that, at least in some cases, high mean levels are indeed caused by a relatively low percentage of cells, meaning that there are still a lot of challenges ahead for efficiently treating these tissues.

One of the features of our cost function is that resource limitations play an important role in shaping the cost landscape. There are indications that cellular NAD levels are limiting, and that a sufficient supply of NAD to mitochondria becomes critical [66, 67, 68, 69]. An increase of intracellular NAD can lead to an increase in oxygen consumption and ATP production [69] indicating that resource limitation may, at least in some cases, be a genuine constraint. Adding various kinds of resources can significantly change mitochondrial basal respiration rate [70, 71, 72].

We found that, under our model, it is more optimal to sense mutants to a lesser extent if the mutants have a lower energy output. In this way, their presence has little influence on wildtype copy number dynamics which then allows the wildtype mtDNA molecules to remain close to their ‘natural’ (and assumed to be close to optimal) levels. This makes a control directly based on the energetic status of the cell a good strategy.

If two different mtDNA types coexist in a cell and one type is more leaky (i.e. less efficient), then this leaky type will tend to produce less energy which could lead to a lower value for *δ.* Cells homoplasmic in the leaky type will then have higher mtDNA copy numbers than cells homoplasmic in the other mtDNA type (see also [73]). Reactive Oxygen Species (ROS) can damage the mitochondrial membrane and increase the amount of leak, making the mitochondria less efficient. It has indeed been suggested that increased ROS production by mutant mtDNA molecules is the reason why cells harbouring these mutations have increased mtDNA copy numbers [74]. It could also be that the increase in mtDNA copy number is due to higher numbers of wildtype mtDNA to compensate for the dysfunctional mutants. Further measurements of mutant as well as wildtype copy number values for different types of mutants, together with their energetic functionality and ROS production rates, are helpful for improving our understanding of the cell’s control mechanisms.

In modelling gene therapies, our fit to endonuclease data yielded a high off-target cutting estimate, ξ = 0.76, which may be a result of the high nuclease concentration in experiments: a lower mtZFN concentration could result in a lower ξ and lead to better results, as was observed experimentally [38]. If the parameter ξ is a function of the treatment strength *I_0_*, the values for I_0,opt_ we found in Figure 5F will be different. Further work on the relationship between *I_0_* and ξ will elucidate more clearly the trade-off between treating with a high strength and cleaving more mutants in the process, and having a high selectivity and therefore low off-target cleavage.

Like any other model, our models have a defined range of applicability. A key baseline assumption was using identical replication and degradation rates for mutants and wildtypes. Various possibilities of distinct rates have been offered in the literature, including faster mutant replication rates [75, 76, 77, 78, 56, 24] and lower mutant degradation rates [79], mainly to explain mutant expansion seen in several tissues. A recent computational model [24] suggested that introducing feedback control on ATP levels, with mutant mtDNAs having a deficient feedback loop, might account for experimental data regarding mutant accumulations. There are also reasons to believe that mutants might be degraded faster, especially if they are dysfunctional, through the mechanisms of quality control [80, 81]. Including such differences, and other features including de novo mutations, degradation control, and cell divisions [82, 52, 42, 83], constitute natural extensions to our theory.

Including these difference in turnover rates, and studying their effects on optimal clinical treatment strategies, provides an interesting future direction of studies. We did not include any cell divisions in our models, meaning that our results are mainly applicable to post-mitotic cells, though cellular division can readily be included (as in, for example, [82] which includes cell divisions during development). Other potential ways in which to develop our models further are including de novo mutations, considering the effect of an extra mutant cost (caused by e.g. ROS production), and including control in the degradation rate. However, the wide set of general insights and biological and therapeutic predictions that emerge even from our simple model suggest that ‘cost-and-control’ modelling of mitochondrial populations is a valuable theoretical approach to reason about these complex and vital systems that remain experimentally hard to address.

## 5 Acknowledgments

We thank Juvid Aryaman and Tom McGrath for useful discussions. The authors declare that there is no conflict of interest regarding the publication of this article.

## Supplementary Information

### S1 Solutions to the system size expansion

We can describe the mtDNA dynamics analytically using a Master Equation:

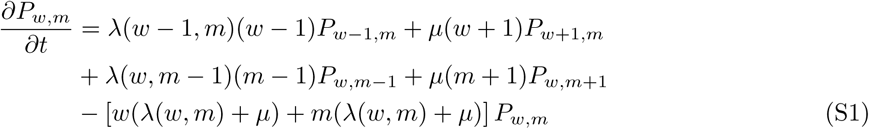

where *P_w_*_,m_*(t)* gives the probability of being in state (*w, m*). In general, this equation will not be analytically solvable and suitable approximation methods are required. We use Van Kampen’s system size expansion to extract a Fokker-Planck equation describing the system, yielding expressions for the means, variances and covariance of *w* and m (table in Figure 1 of the main text). A general master equation can be written in the form

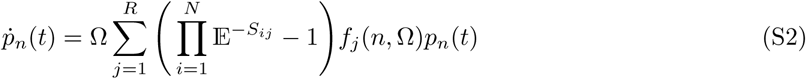

where Ω is the system volume, *R* is the number of reactions involved, *N* is the number of species, *(S)ij* is the stoichiometry matrix, *n =* (*n*_1_, *n*_2_,…,*n_N_)* gives the number of particles of each species, E is a raising and lowering operator^1^, and *p_n_(t)* is the probability distribution of *n* at time *t.* It is often not possible to solve a master equation explicitly, this is only possible in rare cases (the equation *can* be solved analytically for constant or linear rate equations). It is therefore necessary to have approximation methods. The system size expansion, developed by Nico van Kampen, provides a systematic approximation method in the form of an expansion in powers of a small parameter [1]. This parameter, Ω, describes the inverse system size.

To start the construction of the expansion, the state of the system needs to be expressed in terms of a deterministic and stochastic component. Suppose the master equation describes the dynamics of a single species, whose concentration and copy number are denoted by *ϕ*(*t*) and *n.* One expects the distribution of *n* to be centered around Ωϕ*(t)* (with Ω the volume (i.e. size) of the system) and have a width of order 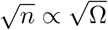 This motivates the starting point of the expansion, which is to write

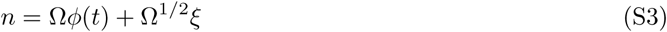

where ξ describes the fluctuations around the deterministic solution *ϕ(t)*.

All the terms in the master equation can now be expressed in the fluctuation variable ξ, according to the following transformations^2^

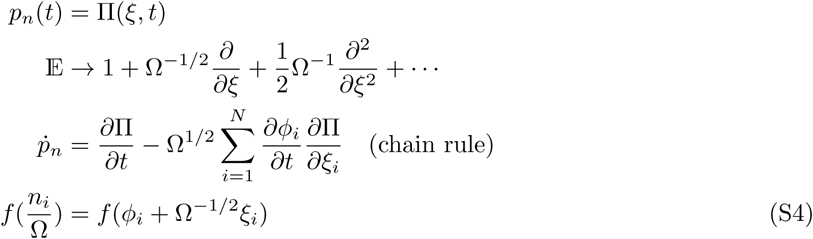

which leads to the new equation

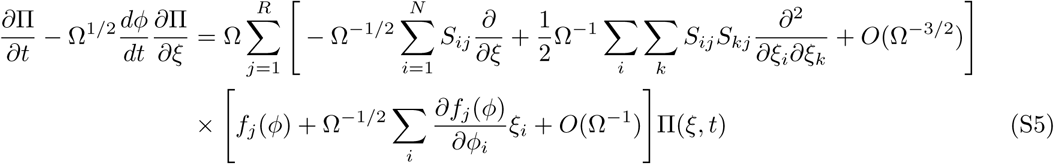

The way to solve this equation is by collecting powers of Ω. The large terms, proportional to Ω*^1/2^*, form the macroscopic rate equations. The terms of order Ω^0^ Form a Fokker-Planck equation, the solution of which is called the Linear Noise Approximation (LNA):

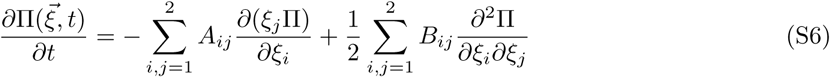

The coefficients *A_ij_* and *B*_ij_ can be found by expanding equation (S5) and are given by

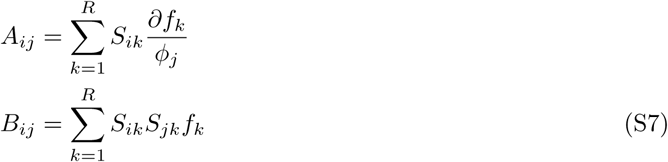

For the system we consider, we have

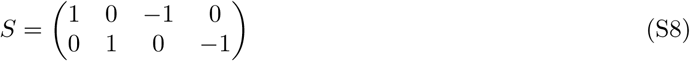

and

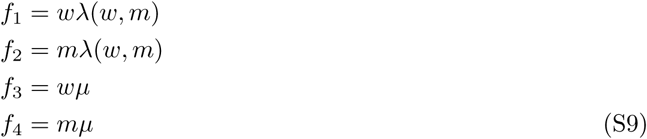

The Fokker-Planck equation can be transformed into a set of coupled ODEs given by

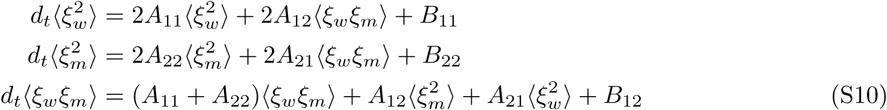

which can be solved to give

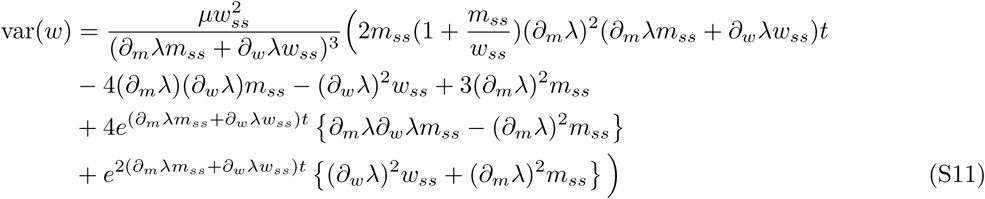

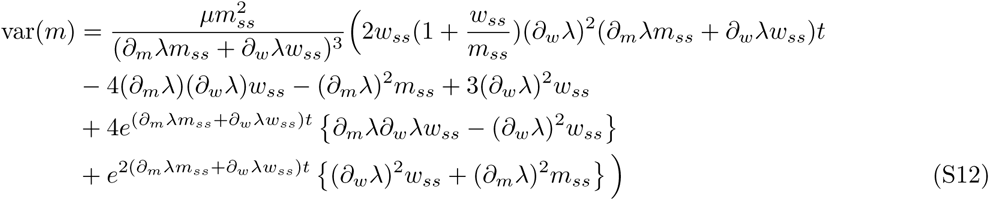

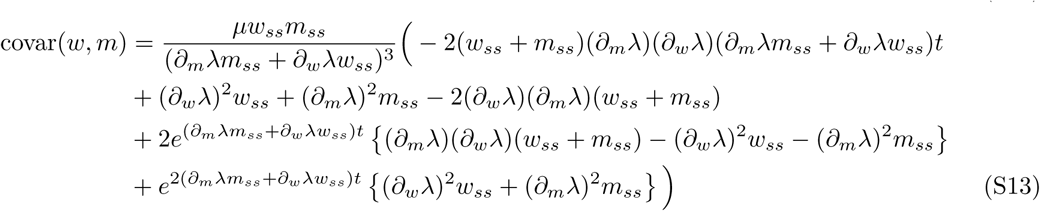

These solutions give the means, variances, and covariance of *w* and m over time according to the Linear Noise Approximation for a general control λ(*w, m*), assuming a trajectory starting in state *(w_ss_, m_ss_).* All the partial derivatives are evaluated at the deterministic steady state (*w_ss_,m_ss_*). The intermediate-time forms of these expressions (when the exponentially decaying terms have died out) are given in Figure 1 in the main text. For very long times (around 1500 days depending on the size the derivative *∂_w_λ* and *∂_m_λ)*, higher order solutions to the System Size Expansion are required to provide a more accurate description of the dynamics.

The heteroplasmy variance can be derived using a Taylor expansion. If only first order are kept, we obtain: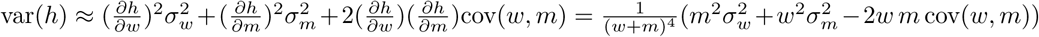.

### S2 Feedback control of a heteroplasmic mtDNA population

#### Steady states of a linear feedback control

When mutant and wildtype mtDNA molecules have identical replication (λ_m_ = *λ_w_*) and degradation (*µ_m_ = µ_w_*) rates, infinitely many steady states exist (i.e. states in which λ = *µ*). These steady states form lines in (*w, m*) space which can be straight (linear feedback control), form segments of ellipses (quadratic feedback control) or take more complicated forms (Figure S1A). Deterministic trajectories will asymptotically approach the steady state line at a specific point; stochastic trajectories can fluctuate along the line thereby changing heteroplasmy.

A linear feedback control *λ*(*w, m*) *= c*_0_ – *c*_1_(*w + δm*) gives rise to a straight line of steady states, the slope of which is determined by *δ.* A small δ means that the steady state line intersects the mutant axis at higher copy number than the wildtype axis, meaning that total copy numbers are higher at *h =* 1 than at *h =* 0. In the extreme case of *δ =* 0, mutant copy numbers can fluctuate off to infinity, though in practice their numbers will be bounded by space restrictions and resource competition. When *δ* > 1 copy numbers will decrease as *h* increases; mutants are now sensed more than wildtypes, which could be caused by e.g. excessive ROS production by mutants (which is then sensed by the cell and incorporated in its feedback function).

**Figure S1:**
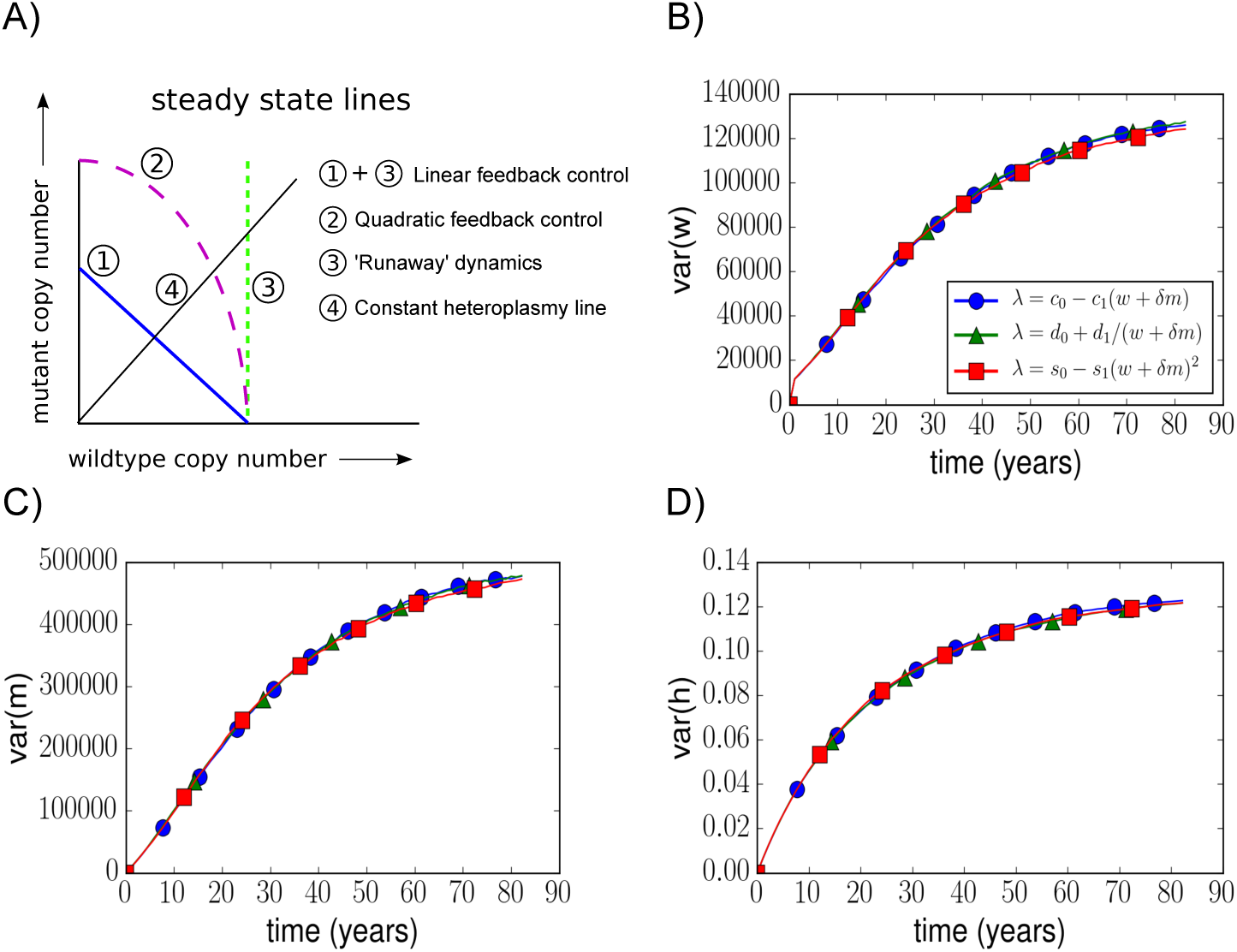
Comparison of different feedback control mechanisms. Related to Figure 1A–C; A) Steady state lines of different kinds of controls are shown in (*w, m*) space. Line ‘3’, a control only dependent on wildtype species, can give rise to ‘runaway dynamics’; mutant copy number can fluctuate off to infinity while wildtype copy number remains stable. B), C), D) Here we show three different controls of the form *λ*(*w* + *δm*) which all have nearly identical wildtype, mutant and heteroplasmy variances. The parameters *c_1_, d_1_, s_1_* are set such that the coefficient of variation of the wildtype distribution in the absence of mutants is given by 0.1. Parameters *c*_0_, *d*_0_, *s*_0_ are set such that the wildtype steady state in the absence of mutants is 1000. Initial conditions *w*_0_ = 919 and *m*_o_ = 162 are used (to give an initial heteroplasmy of *h*_0_= 0.15).

#### Different forms of control can yield similar mtDNA dynamics

We compared three different forms of feedback control, all of the form *λ(w, m) = λ(w* + *δm):* i) *c*_0_ – *c_1_(w* + *δm)*, ii) *d_0_* + *d_1_/ (w* + *δm)* and iii) *s*_0_ – *s*_1_(w + *δm)^2^* with *c*_0_, *c_1_, d_0_, d_1_, s_0_, s_1_ >* 0. Figure 1 in the main text shows that wildtype, mutant and heteroplasmy mean are very similar for each of these controls. Here we illustrate that the wildtype, mutant and heteroplasmy variances are also similar (Figure S1B, C, D).

We merely show that these different controls *can* give rise to similar dynamics by parameterising them such that they do. First, we set the variances of each control to be equal in the absence of mutants. From Table 1E it can be seen that the wildtype variance is now completely specified by the mtDNA degradation rate µ, the steady state copy number *w_ss_* and the control derivative *∂_w_λ*(*w*) evaluated at steady state. The control derivatives at steady state for our three different control mechanisms (as defined above) are given by 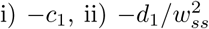 and iii) −2s_1_w_ss_; these expressions were all set to be −7.21 × 10–^6^. This constant value is arbitrary, we chose its value such that at steady state the standard variation of the wildtype distribution is a tenth of its mean value; other choices of this constant will give similar results. We further used *w_ss_* = 1000, which then fixes the values for *c_1_, d_1_* and *s_1_.* To ensure that *w_ss_* forms the steady state, we further set λ(*w_ss_*) = *µ* which determines the values of *c*_0_, *d_0_* and s_0_. Note that it is not surprising that these different controls yield similar dynamics after we have parameterised them to do so; we only want to illustrate these similarities to stress that *which* quantity is being controlled can be more important than *how* it is controlled.

### S3 The relationship between resource consumption and energy production

Mitochondrial respiration is a process by which energy from nutrients is converted into (amongst others) ATP. Part of this process involves the pumping of protons across the inner mitochondrial membrane to create an electrochemical potential across the membrane. Energy is released when protons flow back into the matrix and this energy can be used to create ATP. However, the coupling between proton pumping and ATP synthesis is not perfect and protons can leak through the membrane, reducing the efficiency of respiration. An often measured quantity in experimental studies is the mechanistic P/O ratio [2, 3] which refers to the theoretical maximum amount of ATP (P) produced per oxygen (O) reduced by the respiratory chain. The effective P/O ratio is more physiologically relevant and takes into account leak

[4] In our cost function we need to specify how s_i_, the power supply measured in ATP (including leak) of a mitochondrion of type *i*, depends on *r_i_*, a quantity resembling the resource consumption rate of the mitochondrion. For robustness, we use two different equations for *s*_i_(*r*_i_). The first model is based on measurements in isolated mitochondria which found a linear relationship between *r*_i_ and *s*_i_ (e.g. [5, 6, 4]). We used the data from [5] to fit the parameters of this linear model (Table S1). This data comes from experiments using isolated mitochondria, and it is not clear whether the observations made in isolated mitochondria, without any interactions between the mitochondria and the nucleus or the endoplasmic reticulum, still hold *in vivo.* This is one of the reasons also consider another type of model, as described below.

Our cost function contains a term which penalizes the consumption of resources, meaning that the cheapest state is the one that satisfied demand which the smallest possible resource consumption rate. A linear function *s_i_(r_i_)* then implies that for a given demand, the optimal number of mitochondria to have is the minimum number required to satisfy this demand with the mitochondria respiring as fast as they can. However, a minimal number of mitochondria will make the cell less robust to stochastic fluctuations in both mitochondrial copy numbers and demand. Moreover, mitochondria are known to have large spare capacities [7, 8] indicating that in resting state they do not operate near their limits. We therefore expect that there is some extra cellular cost associated with this ‘maximally respiring’ state, causing it to be non-optimal in resting conditions.

This is why we have chosen to use a second model which describes a saturating relationship between *r_i_* and *s_i_*, as shown in Figure S2. Note that we do not claim that mitochondria become less efficient as they respire faster, we impose the saturating shape merely to effectively assign a higher cost to high respiring states. By imposing this saturation, the variable on the y-axis of Figure S2 can be interpreted an ‘effective energy production’. We will refer to the two models as ‘the linear output model’ and ‘the saturating output model’.

A changing energy production efficiency is not entirely unreasonable, though, because in experiments with isolated mitochondria one usually uses a particular substrate (or a particular combination of substrates) whereas a larger mixture of substrates will be available in the cell, and the relative presence of each substrate may fluctuate over time. The efficiency of respiration depends on the kind of substrate that is used, so it may be possible that at high demand (and high respiration) the substrate of first choice has become limited and another less efficient substrate is used instead. Also, it was suggested that spare capacity can be caused by an increase in substrate entrance in the TCA cycle [9]. This would mean that at high respiration, this high respiration rate is caused by an increase in electron transport chain substrates (e.g. NADH and FADH2). A ‘push’ to the proton pumping complexes instead of a ‘pull’ at the ATP synthase would lead to an increase in the electrochemical gradient across the membrane and therefore an increase in leak. These arguments are speculative but show that the saturation model may not be unreasonable.

The linear and saturating models are given by the equations:

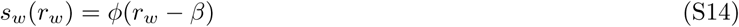

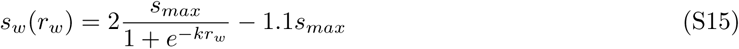

where ϕ can be mapped to the effective P/O ratio (here based on the substrates pyruvate and malate), *β* indicates the respiration rate at zero energy production and therefore specifies the amount of leak, *k* is the rate at which the saturating model saturates, and *s_max_* is an indication of the maximum energy production rate. We have set the parameters *k* and *s_max_* such that the saturating function at first approximately follows the linear function, but then bends away from it at higher respiration rates.

### S4 Parameter values for the cost function

Some of our results will be a consequence of the exact structure of our cost function, and might have been different if another type of cost function was used. We would argue, however, that the main elements in our cost function are quite general: terms involving supply, demand, and resource. We aimed at making our cost function simple, and using biologically interpretable parameters. We do not aim to give a detailed kinetic description of the energetic costs involved, but present a simpler description that allows us to compare distinct strategies relative to each other rather than providing absolute costs. We use our cost function as a tool to characterize cost landscapes and begin to explore optimal control strategies.

In the spirit of ‘back-of-the-envelope’ reasoning in biology [10] we seek plausible and interpretable parameter estimates, using both order-of-magnitude estimations and values found in the literature. The parameters with their default values are summarized in Table S1.

The final goal is to obtain a cost for a state with a certain number of mtDNA molecules; we therefore need to express our cost as a cost ‘per mtDNA molecule’. Because the density of mtDNA molecules within the mitochondrial network seems to be roughly constant [11], we assume every mtDNA molecule is associated to a particular amount of mitochondrial volume which we here define as a ‘mitochondrial unit’. All our parameters refer to these mitochondrial units.

#### S4.1 Mitochondrial energy production and leak: *ϕ* and *β*

In our linear model we assume a linear relationship between mitochondrial ATP production rate and mitochondrial oxygen consumption rate, based on experimental data [5, 4, 6]. In [5, 6] measurements show an almost perfect linear relationship between these two quantities. This relationship can also be determined when the effective P/O ratio (the amount of ATP produced per oxygen consumed) is known as a function of the oxygen consumption rate; this also leads to a linear relationship between ATP synthesis rate and oxygen consumption rate [4].

The parameters *ϕ* and *β* correspond to the slope and intercept of the linear function, respectively. In [5] this slope was measured to be 2.03 ± 0.13 for isolated pectoralis muscle cell mitochondria in the presence of pyruvate and malate; we have decided to use ϕ = 2, mainly based on these experiments.

The value of *β* is an indication of the ‘leakiness’ of the mitochondrion: it represents the rate of oxygen consumption that is required to balance the leakage of protons across the membrane in order to maintain the mitochondrial membrane potential. To obtain a consistent value for β we also use the data presented in [5]. Their measurements find that *β* is about a tenth of the maximum respiration rate. This maximum respiration rate is obtained by adding high (unlimited) concentrations of ADP; the state of the cell in these conditions is known as state 3_ADP_. Because state 3_ADP_ does not necessarily correspond to *in vivo* conditions, we define the respiration rate in this state as *r_max_*_,t_: the maximum ‘theoretical’ respiration rate. We will fix *r_max,t_ =* 1 and use this to scale our other parameters. This means that our parameter value for *β* is *β* = 0.1.

We stress that though we have based our parameter values here on a specific study, changes in their values will not affect the qualitative structure of our cost function but merely changes the slope and intercept of the linear output function defined in equation (S14).

#### S4.2 Resource and supply and maintenance cost: *r_max_,r_n_* and *s_n_*

A cell *in vivo* is unlikely to experience the high concentrations of ADP present in state 3_ADP_. We therefore set our parameter *r_max_*, the physiological maximum respiration rate, to be slightly below *r_max_*_,*t*_*:r_max_ = 0.95r_max,t_ =* 0.95.

We introduce *r_n_* and *s_n_* as the normal respiration rate and ATP production rate which are present in resting conditions, respectively. Their values will be used to derive other parameter values. Mitochondria have spare capacity, i.e. in normal unstressed conditions they use only part of their maximal oxygen consumption rate (OCR). The amount of spare capacity is usually measured as the fold-change in OCR that occurs after adding FCCP to cells, a mitochondrial uncoupler. Several measurements of the fold-change in OCR are: in the range (2–4)-fold [12], 1.4- to 2.5-fold [9], about 2-fold [7] and about 2.5-fold [8]. The maximum respiration rate when adding FCCP is known as State 3_FCCP_, and is higher than State 3_ADP_ (see e.g. [13]). We interpret the spare capacity as the ratio State 3_FCCP_/r_n_, meaning that the ratio *r_max_/r_n_* is lower than this. We have decided to take *r_max_/r_n_ =* 1.5, meaning that *r_n_ = r_max_*/1.5 ≈ 0.63. The value for *s_n_* is now simply *s_n_ = s_w_*(*r_n_*) ≈ 1.1.

### S4.3 Maintaining, building and degrading: ρ_i_, ρ_2_ and ρ*_3_*

The model organism with the most quantitative data on mitochondrial energy budgets is budding yeast, Saccharomyces cerevisiae. Reasoning that the scales of biophysical costs of mitochondria are likely comparable across eukaryotes, we first draw from this literature to motivate order-of-magnitude estimates for a range of essential mitochondrial processes. We will later construct parallel estimates using other organisms.

We first focus on estimating the mitochondrial building cost (in ATP) ρ_2_. We provide three distinct estimations and combine them to obtain our final estimate.

For the first estimation we use a list of mitochondrial proteins in yeast [14], and obtain information on turnover rates, abundance (per yeast cell), and lengths (amino acid length) of these proteins by using the Saccharomyces Genome Database [15]. We end up with a list of about 200 mitochondrial proteins in yeast *S. cerevisiae.* We incorporate the observation that it takes about 5.2 ATP molecules to elongate a growing peptide chain by adding an amino acid [16], which means that the total synthesis cost of the mitochondrial proteins included in our list is given by 5.2 Σ*_i_* length_*i*_abundance_*i*_ ≈ 2 × 10^10^ ATP per yeast cell. The known number of mitochondrial proteins in *S. cerevisiae* is on the order of 1000 [17]; we will therefore assume that the protein synthesis cost obtained from our protein list corresponds to roughly a fifth of the total mitochondrial protein synthesis cost. We might expect that the proteins best known [14] are the more abundant ones, meaning that our final cost is likely to be an overestimate. We also assume that all of the mitochondrially associated proteins are used exclusively for mitochondrial function. In *E. coli*, the mitochondrial protein synthesis cost represents ~50% of the total mitochondrial synthesis cost (two other major contributors are phospholipid synthesis and RNA synthesis)[16]; we will make the assumption that this observation in *E. coli* holds in mitochondria as well. This brings the total mitochondrial building cost in a single yeast cell to be about 1.9 × 10^11^ ATP. Assuming 50–100 mtDNA molecules per yeast *S. cerevisiae* cell [18], the building cost associated to a single mtDNA molecule (and therefore the building cost of a mitochondrial unit) is given by (2 – 4) × 10^9^ ATP.

For our second estimation we use the total protein weight of a single mitochondrion which was measured to be about 3 × 10^−10^ mg in rat liver [19], as well as the typical weight of a single protein which is about 5 × 10^-17^ mg [16]. This means that a mitochondrion contains about 6 × 10^6^ proteins. Using that the typical length of a protein is 300 amino acids [20, 21, 16] together with the 5.2 ATP cost of adding amino acids and the estimation that protein costs represent 50% of the entire building cost, the mitochondrial building cost is estimated to be 2 × 10^10^ ATP. Note that this cost does not necessarily represent our mitochondrial unit because it is unknown how many mtDNAs a ‘mitochondrion’ corresponded to when measuring its weight in [19].

The third estimation is based on the building cost of an *E. coli*, which is about 10^10^ ATP [16]. Keeping in mind that we want the building cost of a fraction of mitochondrial volume corresponding to a single mtDNA molecule, we need to convert the building cost of an *E. coli* (which has a volume of about 1 μm^3^) to represent our mitochondrial unit. It was estimated that the total mitochondrial network length in yeast *S. cerevisiae* is about 25 μm with a total mtDNA copy number of 50–100 [18]. Assuming that the mitochondria form tubules with a constant diameter of 300 nm [22] gives a total mitochondrial volume of about 1.8 μm^3^; another total mitochondrial volume estimate in yeast *S. cerevisiae* is 1.5 μm^3^ [22]. Assuming uniform distributions for the mitochondrial volume (1.5–1.8 μm^3^) and mtDNA copy numbers (50–100) leads to a volume of (2.3 ± 0.5) x 10^−2^μm^3^ per mitochondrial unit. This means that rescaling the *E. coli* building cost gives us an estimate of (2.3 ± 0.5) × 10^−2^ ⋅ 10^10^ = (2.3 ± 0.5) × 10^8^ ATP to build a single mitochondrial unit. Note that this may represent an overestimation because *E. coli* is a unicellular organism, whereas the mitochondrion is an organelle which cannot survive in isolation [23].

While there are differences in these estimates, arising both from uncertainty and different quantitative lines of reasoning, they together give an overall scale for mitochondrial building cost of around 10^9^ ATP. Because in our model we only need a rough estimate of the mitochondrial building cost, we use *ρ_2_ =* 10^9^ ATP.

We interpret the maintenance cost, denoted by ρ_1_, as the cost in molecules ATP/s corresponding to, for example, maintaining the mitochondrial lipid membranes, importing/exporting proteins, and synthesizing new proteins. To obtain an estimation of ρ_1_, we again use the Saccharomyces Genome Database [15]. We can calculate the cost of continuously turning over the ~ 200 proteins in our list (obtained from [14]), leading to 5.2 Σ*_i_* length_*i*_ abundance*_i_* (degradation rate)*_i_* ≈ 6 × 10^5^ ATP/s per yeast cell. In other words, the maintenance cost per second of our set of mitochondrial proteins is about five orders of magnitude less than their synthesis cost. We therefore assume ρ_1_ *= 10^−5^ρ_2_.*

ρ_3_ is the most challenging parameter to estimate, as the process of mitochondrial degradation remains poorly characterised. Protein production and biosynthesis costs form the bulk of mitochondrial production requirements, and from cell-wide studies on energy budgets are among the most considerable demands in cell biology. We therefore assume that degradation has lower energy requirements than production, and set its upper limit at ρ_3_ *=* 0.1ρ_2_. Lowering ρ_3_ further has little impact on the outcomes of our model.

#### S4.4 Demand and resource availability: *D* and *R*

We want to model two kinds of cells: low copy number cells (1000 wildtype mtDNAs in resting state) and high copy number cells (5000 wildtype mtDNAs in resting state). Denoting this desired number of ‘normal’ mitochondria by *w_n_*, we obtain

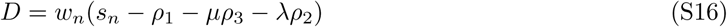

where *µ* and λ are the degradation and replication rates (per second). This equation says that the overall net output of *w_n_* mitochondria exactly satisfies the demand. Using *w_n_ =* 1000 and *w_n_ =* 5000 leads to *D* ≈ 1055 and *D ≈* 5275 ATP/s.

The parameter *R* denotes the maximum rate at which resource can be consumed by all of the mitochondria together and represents a cellular resource availability. In normal resting state the total resource that is consumed is *w_n_r_n_*, and the resource consumed when these mitochondria respire as fast as they can is *w_n_r_max_.* We then assuming this maximal respiration rate is achieved by using all of the available resources, i.e. *R ≈ w_n_r_max_.* It may be, however, that a state of maximum respiration can only be maintained for a short time, and in our cost function we want to describe the ‘steady state cost’ for different states. We therefore assume that *R* < *w_n_r_max_ (R* now denotes the maximal respiration rate that can be maintained for longer periods of time). We have assumed *R = 0.8w_n_r_max_* leading to *R =* 760 and *R =* 3800 for *w_n_ =* 1000 and *w_n_ =* 5000, respectively.

We note that we base the values for the parameters *s_n_ = s_w_(r_n_)* (and therefore the other parameters) on the linear model, with the idea that these are ‘intrinsic’ mitochondrial parameters, because we have data for this linear model. The saturating model keeps the same intrinsic parameters but is different because of influences from the cell itself.

#### S4.5 Saturating model parameters: *s_max_* and *k*

We have chosen the parameters of the saturating model described in equation (S15) to approximately match the linear model for low respiration and reach a lower final ATP production rate for higher respiration. The values we used are *s_max_ =* 1.54 and *k =* 3.0.

#### S4.6 The cost of resource consumption *α*

The value of α, i.e. the scaling parameter that appears in the cost function given by

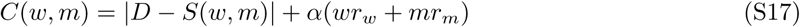

is hard to determine. Its value describes the cost of a unit of resource consumption relative to the cost of a unit of ‘energy deficiency’ (the cost of *S(w, m*) being one energy-unit below *D).* We estimated that a penalty for resource consumption usage should be about an order-of-magnitude less than the penalty for not-satisfying demand, and have therefore decided to assume *α =* 0.1. We note that the value of *α* has no influence on the shape of the demand-satisfying region, it changes the relative costs within (and outside of) the region.

#### S4.7 Mutant parameters: *ϵ*_1_, *ϵ*_2_

In the main text we vary the parameter *ϵ_1_*, describing the resource uptake rate of mutants relative to wildtypes. Additionaly, mutants can be less efficient than wildtypes, producing less energy per resource consumed; we denote this lower mutant efficiency by *ϵ*_2_ ∈ [0,1]. Because the number of protons that are pumped across the mitochondrial inner membrane by the electron transport chain complexes for every unit of resource (NADH) that is consumed is fixed, a lower *ϵ*_2_ would have to mean that either i) the mutation has increased proton leak (or other ways of depolarising the membrane), or ii) the mutation has made the ATP synthase dysfunctional. The value of *ϵ*_2_ can be related to the P/O ratio of the mutants relative to that of the wildtypes. Most mtDNA mutations, however, affect the electron transport chain complexes themselves and are therefore likely to reduce the flow of resources through the chain (which would mean a low value for *ϵ*_1_). This is why we assume *ϵ*_2_ = 1 in our main model and only vary the parameter 0 < *ϵ*_1_ < 1. For completion, here in the SI we provide a heatmap showing the cost in *(w, m*) space for various values of *ϵ*_2_ (Figure S3).

### S4.8 Parameter units

We can relate our parameter values to actual values, e.g. expressing our demand *D* in ATP/s. As an example of an ATP demand we use the ATP production rate in unstressed human skin fibroblasts. Assuming that these healthy cells satisfy their demand, their net ATP production rate should equal their ATP demand. The rate of ATP production in skin fibroblasts was estimated to be about 10^9^ ATP/s, the large majority of which is supplied by mitochondria [24]. The number of mtDNAs in healthy human skin fibroblasts was measured to be roughly in the range 2400–5200 [25] (the variation in copy number was partly due to variation in ages of the individuals), and we will use the value 4000 as an estimation. Using *w_n_ =* 4000 we obtain

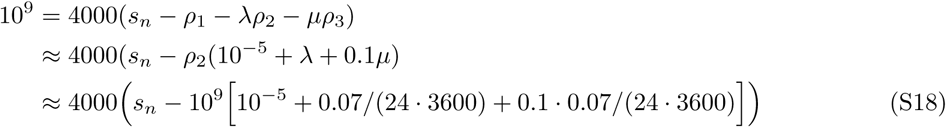

Here we used an mtDNA half-life (T_1/2_) of 10 days, giving a degradation rate ln(2)/10 ≈ 0.07 day^−1^, and assumed that the cells are in steady state with λ = *µ.* This leads to *s_n_ ≈* 2.6 x 10^5^ ATP/s meaning that, considering we used *s_n_ ≈* 1.1, our parameters *s_n_,D* are expressed in units of about 2.6 x 10^5^ ATP/s. This means that, in our units, the parameter values we use for ρ_1_*, ρ_2_* and ρ_3_ are ρ_1_ *≈* 0.04, ρ_2_ *≈* 3828, and ρ3 *≈* 383.

**Table S1:** Parameters used in our cost function with their descriptions

| Par. | Description | default value |
| --- | --- | --- |
| $D$ | Mitochondrial energy demand in ATP/s. | 5110.7 (high copy number)<br>1022.1 (low copy number) |
| $R$ | Maximum rate of resource supplied by the cell to be used by the mitochondria. We use the term ‘resource’ as an amalgamation of different mitochondrial resources, e.g. NAD(H), pyruvate, lactate, succinate, ADP, $P_i$, and oxygen. | 3800 (high copy number)<br>760 (low copy number) |
| $\phi$ | $\phi$ is related to the effective P/O ratio of a wildtype mtDNA molecule, it represents the slope of the linear relationship between $r_w$ and $s_w$ given in equation (S14) | 2.0 |
| $\beta$ | The fraction of the maximum respiration rate that consists of proton leak, in ‘resource’ per second | 0.1 |
| $k$ | Parameter describing the saturation of the saturating model. | 3 |
| $s_{max}$ | An indication of the aximum energy supplied by a wildtype mtDNA molecule in the linear model in ATP/s. | 1.54 |
| $r_{max}$ | Maximum rate of resource uptake by an mtDNA molecule (i.e. by a mitochondrion containing the mtDNA molecule). This can be interpreted as the maximum flow through the respiratory chain. | 0.95 |
| $\rho_1$ | The mitochondrial maintenance cost | 0.04 ($10^4$ ATP/s) |
| $\rho_2$ | The mitochondrial building cost | 3828 ($10^9$ ATP) |
| $\rho_3$ | The mitochondrial degradation cost | 383 ($10^8$ ATP) |
| $\epsilon_1$ | The mutant energy production efficiency (the energy produced per unit of resource consumed) relative to that of wildtype mtDNA molecules. A low value of $\epsilon_1$ can be caused by a high proton leak or a deficient ATP synthase. | free parameter, $\epsilon_1 \in [0, 1]$ |
| $\epsilon_2$ | The mutant resource uptake rate relative to that of wildtype mtDNA molecules. We will mainly use $\epsilon_1 = 1$ and $\epsilon_2 < 1$ because mtDNA mutations usually affect the proton pumping electron transport chain complexes. A defect in e.g. complex I will reduce its activity and therefore also the rate of consumption of NADH. | free parameter, $\epsilon_2 \in [0, 1]$ |
| $\alpha$ | Scaling parameter in the cost function in front of resource consumption term. | 0.1 |
| $w_{opt}$ | The cheapest value for $w$ when $m = 0$, using all of the above parameter values. We have four values for $w_{opt}$, corresponding to each of the four systems we study. | 1524 (saturating low)<br>7616 (saturating high)<br>638 (linear low)<br>3129 (linear high) |

### S5 Cost function outputs

Figure S2A shows the behaviour of equations (S14) – (S15) for various parameter values. Note that none of the lines in the figure crosses the origin, because even when no ATP is created, respiration is required to maintain the gradient which would otherwise be lost due to proton leak. Figure S2B–E shows how resource consumption rate and cost change as the number of mtDNAs changes. Three regimes can be distinguished: 1) there are too few mitochondria present to satisfy demand and they use their maximum possible resource uptake to get their energy production rate as close to D as possible; 2) demand can be satisfied and the cost now only depends on the amount of resource that is used; 3) resource has become limiting and demand cannot be satisfied any more. The main difference between the two models is that in the linear model, when demand is satisfied, the cheapest state is the one with the smallest mtDNA copy number possible whereas the saturating model is cheapest at a higher copy number. This is because mitochondria in the saturating model becomes more efficient as less resources are being consumed.

**Figure S2:**
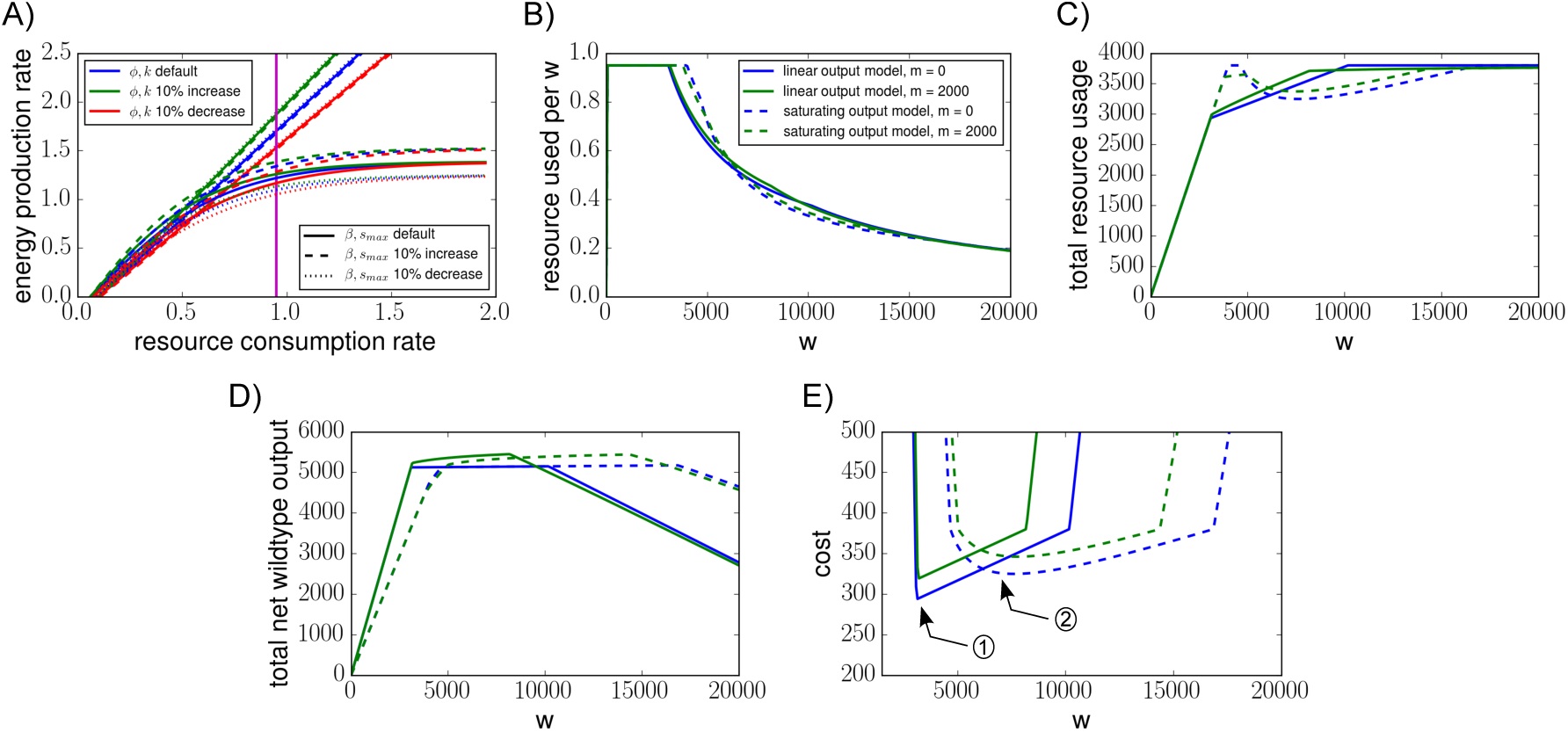
Relationship between resource consumption and energy output. **A)** Here we show the energy production rate of a single wildtype mitochondrion as a function of its resource consumption rate, as given by equations (S14) – (S15). For the linear model (corresponding to the linear lines in the figure) the parameters ϕ and *β* are changed by 10%, for the saturating model (corresponding to the saturating curves in the figure) we vary *s_max_* and *k.* The magenta line indicates the value of *r_max_.* Our default parameters for the linear model are based on data given in [5] **B)** As *w* increases, the demand is shared between more mitochondria and each individual mitochondrion can afford to consume resources at a lower rate (the same figure legend applies for figures C, D and E). C) The total resource consumption does increase as *w* increases because the mitochondria need to consume a non-zero amount of resources to produce a net energy output and each mitochondrion comes with a maintenance cost. **D)** The total energy produced by wildtype mitochondria increases when mutants are present because the mutants have a net energy deficit. **E)** When demand is satisfied, the cost increases with *w* in the linear model, meaning that the minimum cost occurs when mitochondrial copy number attains the minimum number required to satisfy demand (1). In contrast, for the saturating model the cost decreases at first because as the individual resource consumption drops, the energy production efficiency increases. Eventually the cost increases again for similar reasons as in the linear model; minimum cost now occurs when mitochondria are working most efficiently (2). Parameters *ϵ*_1_ = 0.1 and *ϵ*_2_ = 1.0 were used.

In the main text we saw that, in the saturating model, intermediate heteroplasmy values seem to be more expensive than high or low heteroplasmy values. Here we illustrate that in high heteroplasmy regions it is more efficient to increase heteroplasmy even more, whereas in low h conditions it is more efficient to decrease h; this automatically implies there exists some intermediate heteroplasmy value that is least efficient. Figures S3A, B show the amount of resource consumed by the individual wildtype and mutant mitochondria in four different states. All states have identical total copy numbers (*w* + *m* = 10^4^) but different heteroplasmy values (*h* = 0.1, 0.3, 0.7 and 0.9). All four cases have identical total outputs (equal to the demand). When heteroplasmy increases, the individual resource consumption rates *r_w_* and *r_m_* both increase to compensate for the higher mutant copy number; this is true both in a low-*h* region (*h* increases from 0.1 to 0.3, Figure S3A) and a high-*h* region (*h* increases from 0.7 to 0.9, Figure S3B). However, the total resource consumption rate does not necessarily increase because the increase in *h* has caused a number of wildtype mitochondria to become mutants, thereby decreasing their resource usage. Computing the values of the resource consumption rates for the states with *h* = 0.1 and 0.3, while referring to Figure S3A, gives:

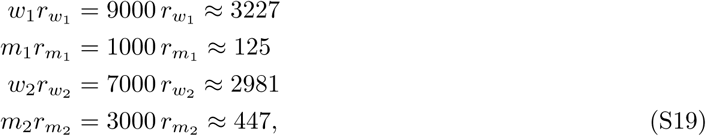

while the states *h* = 0.7 and 0.9 (referring to Figure S3B) give

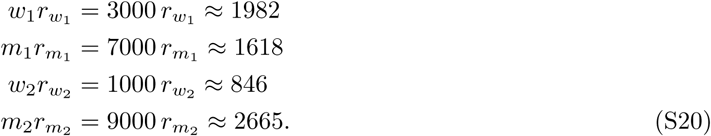

Comparing the states with *h* = 0.1 and *h* = 0.3, the *total* rates of resource usage are 3352 and 3428 respectively; the lower heteroplasmy state is more efficient. However, when heteroplasmies are higher, the high heteroplasmy state (0.9 rather than 0.7) is most efficient. This effect is due to the non-linearity of Equation S15.

**Figure S3:**
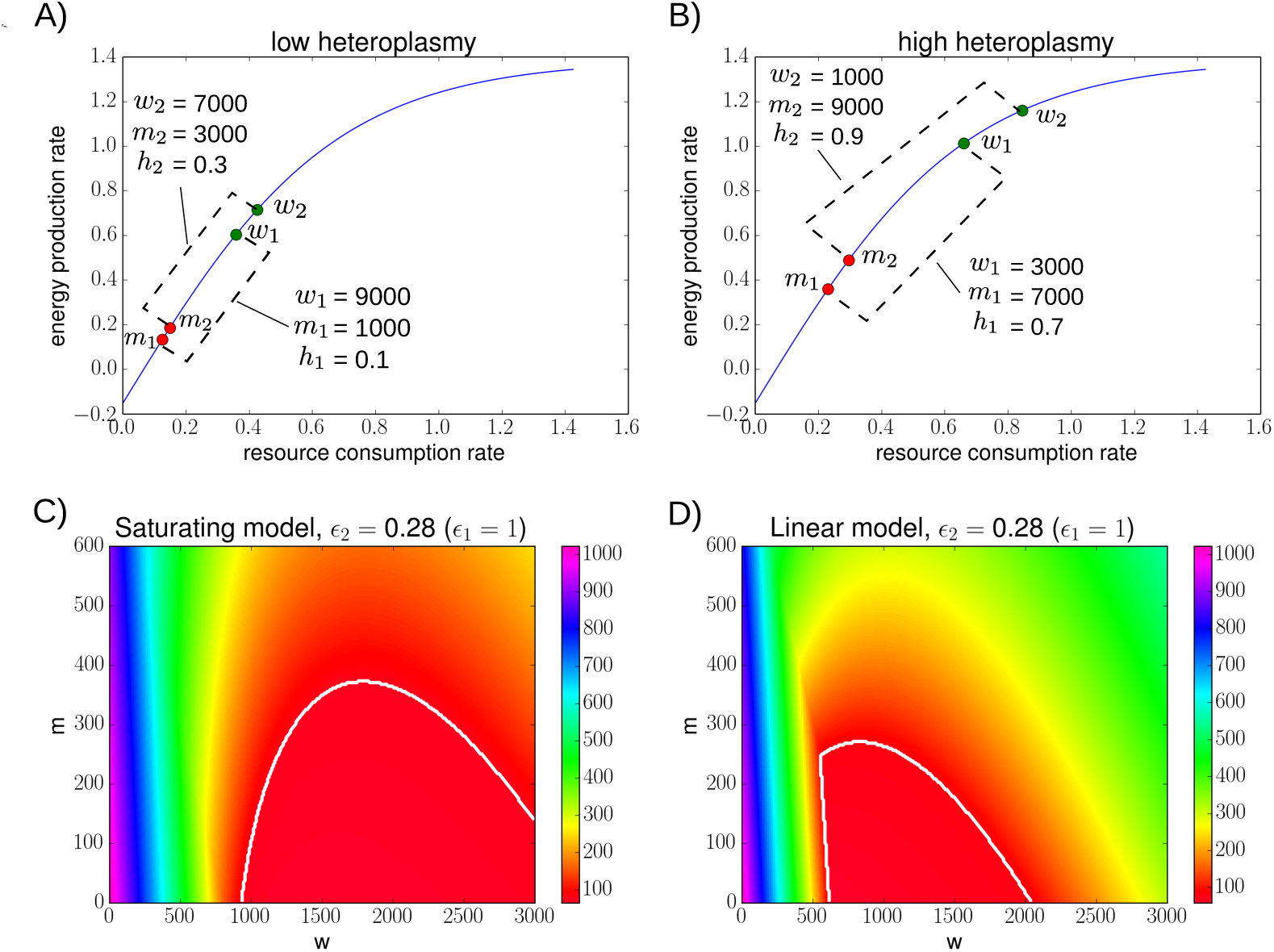
Intermediate h values require more resources to satisfy demand, but only if mutants consume less resources. A) The resource consumption rates and energy production rates of wildtypes and mutants are shown for two states: *(w_1_,m_1_,h_1_)* = (9000,1000,0.1) and *(w_2_*,m*_2_,h_2_)* = (7000,3000,0.3). In both cases, the total energy output is equal to the demand. When heteroplasmy is higher *(h* = 0.3), the individual resource consumption rates are higher in order to maintain a constant total energy output. Overall, the state with *h* = 0.1 uses the least resources (Equations S19). *ϵ_1_* = 0.35 was used. B) This figure is similar to Figure (D) but now the two states *(w1,m1,h1)* = (3000,7000,0.7) and *(w_2_*, m*_2_, h_2_)* = (1000, 9000, 0.9) are compared. The state with *h* = 0.9 uses the least resources (Equations (S20)). C) + D)) Similar to Figure 2 in the main text, these figures show the cost values in *(w, m)* space, but now as a function of 62 (mutant efficiency) instead of e_1_. This time we show the cost in the entire space. The white lines show the region in which demand is satisfied for our default parameter values. Because mutants consume the same amount of resource as wildtypes (e_1_ = 1), resource becomes limiting at relatively low values of *m* compared to when *ϵ_1_* < 1. Note that intermediate heteroplasmies are not less efficient here.

Finally, Figures S3C, D show the cost function as a heatmap in (*w*, *m)* space. This figure is similar to Figure 2 in the main text but now we have fixed *ϵ*_1_ = 1 and varied *ϵ*_2_. Mutants are now less tolerated because they consume just as many resources as wildtypes, but still produce less output. It is now not the case that intermediate heteroplasmies are less efficient; *intermediate heteroplasmies are only less efficient when *ϵ*_1_ < 1 (mutants consume less resource) and when a saturating output model* (Figure S2) is used. When *ϵ*_1_,*ϵ*_2_ < 1, it is possible for intermediate heteroplasmies to be less efficient but the smaller the value of *ϵ*_2_, the smaller the range of values of *ϵ*_1_ for which this is true; therefore, the effect of intermediate heteroplasmies being less efficient will be most easily observed when the mutant efficiency is close to that of the wildtypes.

### S6 Comparison of the cost of different control mechanisms

#### High costs for not sensing mutants are caused by increases in mean mutant copy number

In Section 2.3 of the main text we introduced four different feedback controls and compared their cost. These controls are summarized in Table S2 and will be referred to by their labels as provided in the table. In order to compare the controls we imposed two constraints: we demand that, in the absence of mutants, the controls have i) the same deterministic steady state, which is set to be *w_opt_;* and ii) the same steady state variance. It seems a reasonable assumption that any control would be optimal or near optimal in a ‘healthy’ mutant-free state. The goal is to have the controls behaving similarly when *m* = 0 and observe how the dynamics change in the presence of mutants.

As mentioned before (Section S2), when *m* = 0, the wildtype variance is completely specified by the mtDNA degradation rate *μ*, the steady state copy number *w_opt_* and the control derivative *∂_w_λ(w)* evaluated at steady state. We here determine the magnitude of *∂_w_λ(w)* by fixing the parameter α*_r_* in the ‘relaxed replication model’ [26, 27]; its value was suggested to lie between 5 and 17 and here we have used α*_r_* = 10. All controls are set to have equal values of *∂_w_λ(w)*, and *λ(w_opt_) = μ* (with *m* = 0) is used to provide identical steady states (Table S2). Any free parameters left (*δ* in Control D and γ in control A) are optimized with respect to our cost function; both these optimized parameters specify the relative mutant contribution to the control.

Figure S4 shows the means and variances of the wildtypes, mutants and cost up to ~ 82 years resulting from stochastic simulations. The four controls all started in steady state with *h_0_ =* 0.15. We see that the increase in cost for control B is mainly caused by the increase in mean mutant copy number.

**Figure S4:**
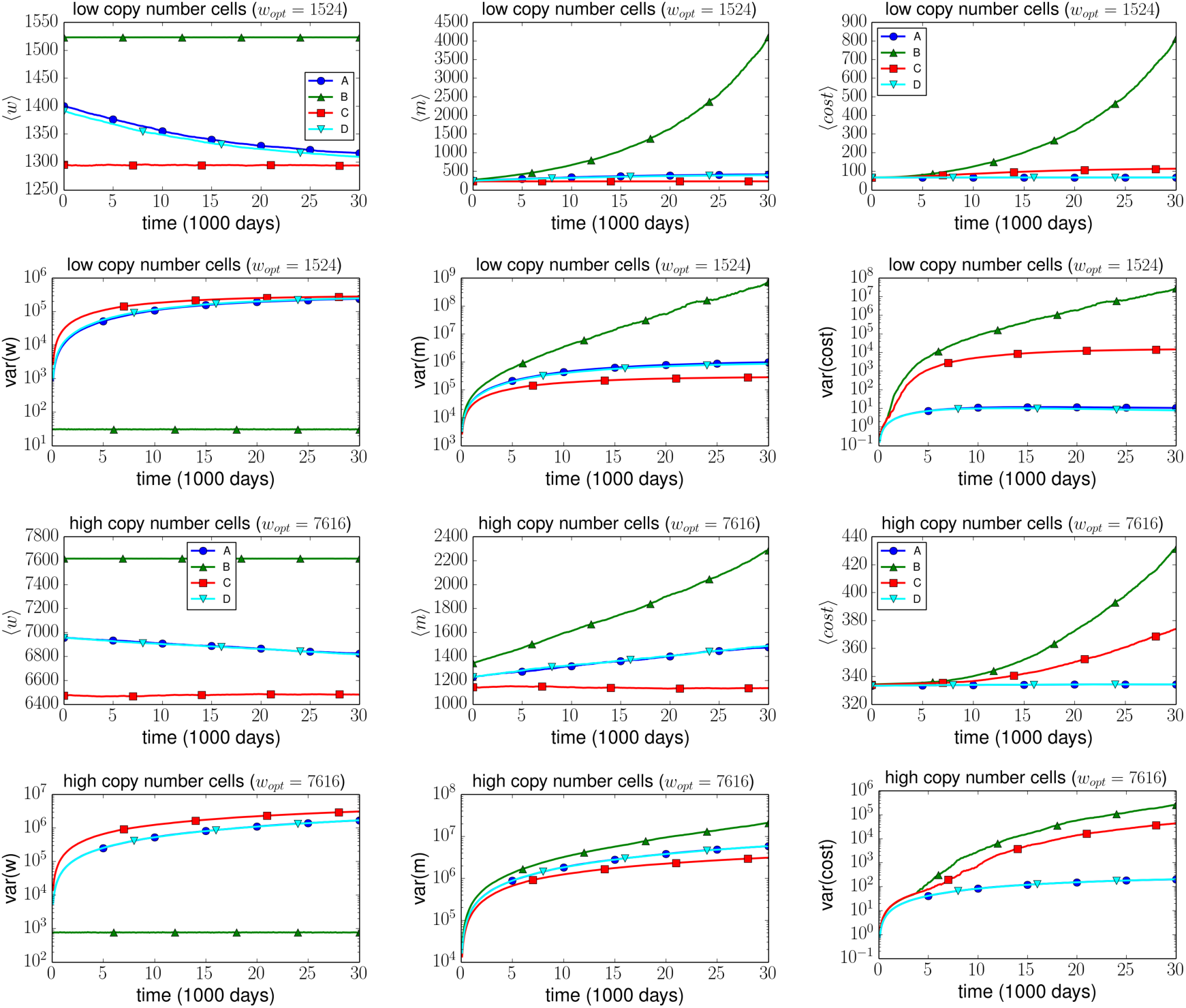
Wildtype, mutant and cost dynamics for four different control strategies. Dynamics are shown for the four controls A, B, C and D given in TableTab: Section 3 tab SI. Again, we see that the effects of the control are more noticeable in low copy number cells. Parameter are set as given in Table S2; the degradation rate used corresponds to a half-life of 10 days (*µ* ≈ = 0.07). Values for *w_opt_* are those for the saturating output model at low and high copy number. The free parameters in control A and D (γ and *δ)* were optimized over initial conditions in the range *h ∈* [0,0.2]. For the optimization the default cost function parameters were used as well as *ϵ_1_* = 0.3

**Table S2:**
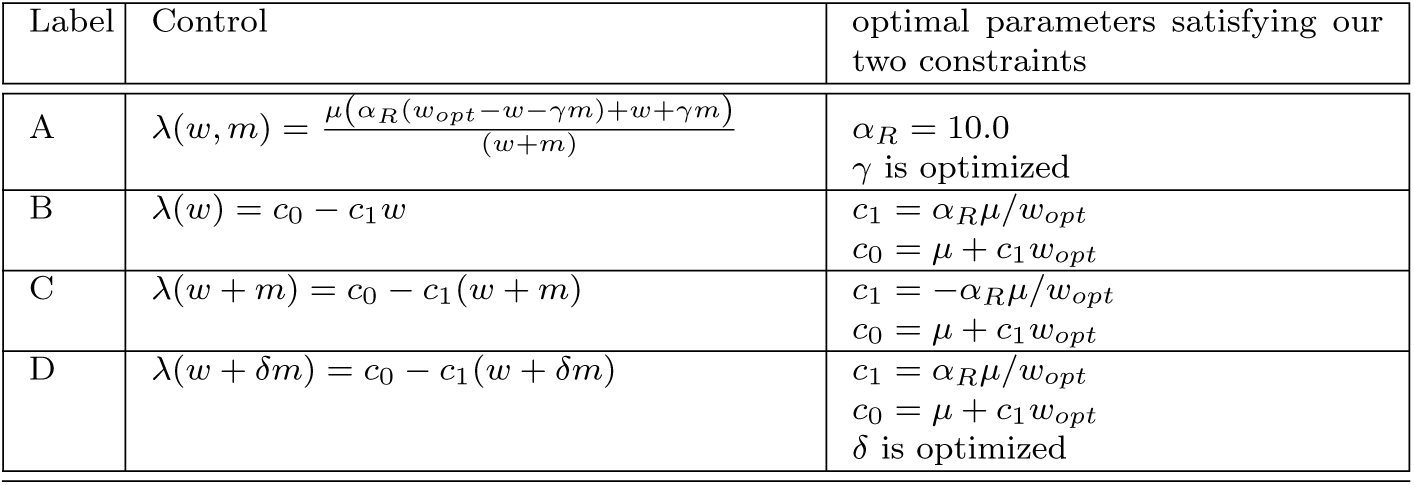
Four controls considered with their parameter values. Two parameters of each control are set by the two constraints we impose. The parameter α_*R*_ was proposed to lie in the range 5–17 [26] and here we used α_*R*_ = 10. The values for *δ* and γ are found by optimizing our cost function over the steady states corresponding to our initial conditions. We used 50 initial conditions equally spread over the range *h_0_ ∈* [0,0.2]. The two values used for *w_opt_* are 1524 and 7616 (Table S1).

| Label | Control | optimal parameters satisfying our two constraints |
| --- | --- | --- |
| A | $\lambda(w, m) = \frac{\mu(\alpha_R(w_{opt} - w - \gamma m) + w + \gamma m)}{(w + m)}$ | $\alpha_R = 10.0$<br>$\gamma$ is optimized |
| B | $\lambda(w) = c_0 - c_1 w$ | $c_1 = \alpha_R \mu / w_{opt}$<br>$c_0 = \mu + c_1 w_{opt}$ |
| C | $\lambda(w + m) = c_0 - c_1 (w + m)$ | $c_1 = -\alpha_R \mu / w_{opt}$<br>$c_0 = \mu + c_1 w_{opt}$ |
| D | $\lambda(w + \delta m) = c_0 - c_1 (w + \delta m)$ | $c_1 = \alpha_R \mu / w_{opt}$<br>$c_0 = \mu + c_1 w_{opt}$<br>$\delta$ is optimized |

#### Optimal linear controls for various mutant pathologies

Figure S5 shows the optimal values for δ in the linear feedback control λ(*w,m*) = *c*_0_ − *c*_1_(*w* +*δm*) as a function of *ϵ_1_*. Stochastic simulations starting in steady state at either *h_0_* = 0.1 or *h_0_* = 0.8 were performed for 10^4^ days. The mean integrated cost over these 10^4^ days was evaluated for different values of *δ*, and the optimal *δ* values are shown. This was done for both the linear and the saturating model. The general trend is, as was shown for *T* = 100 in the main text, that the lower *ϵ_1_* the lower the optimal *δ.*

**Figure S5:**
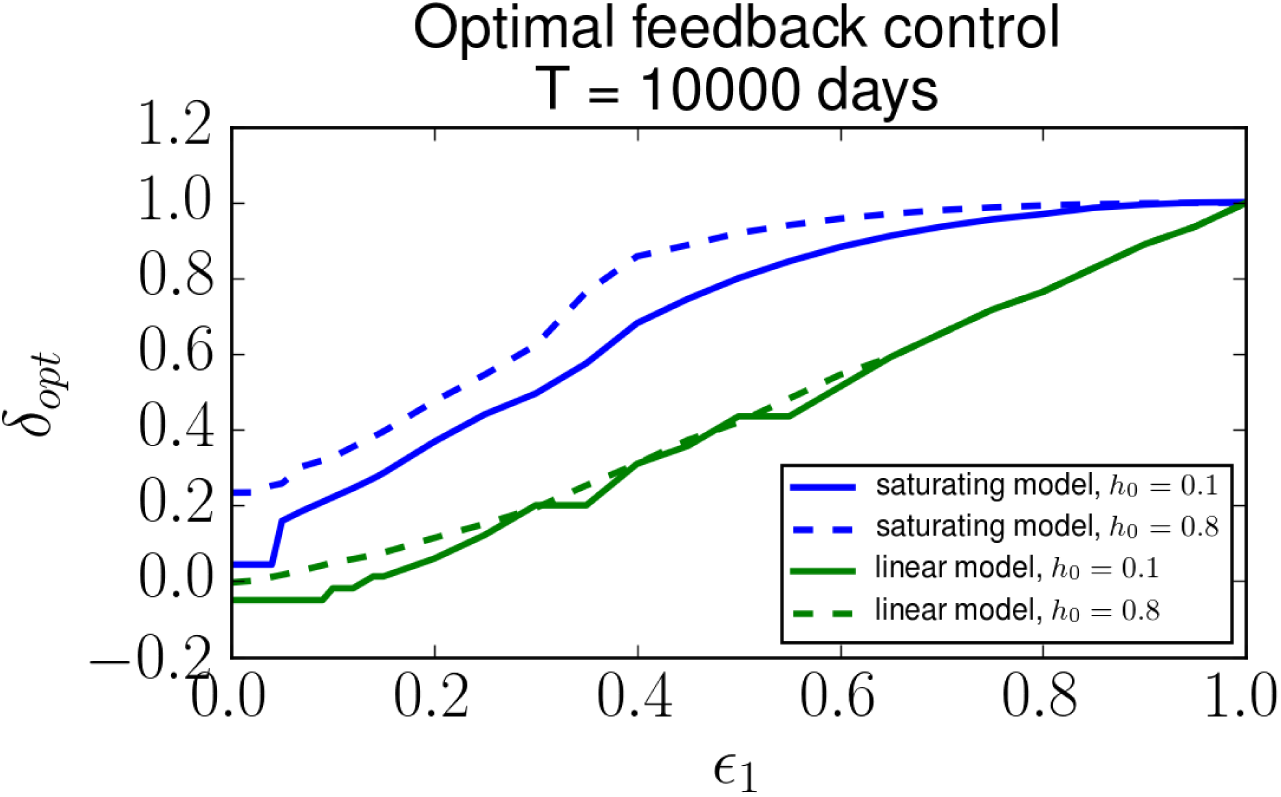
At long times and high heteroplasmies, energy sensing control becomes suboptimal. Related to Figure 3C; The optimal value of *δ* in a linear feedback control is shown as a function of *ϵ_1_*. Here we used *T* = 10000 (optimization time) and low copy numbers for both the linear and saturating model. The solid and dashed lines correspond to trajectories starting at *h_0_* = 0.1 and *h_0_* = 0.8, respectively. The less resources the mutants consume (and the less output they therefore produce) the lower their optimal contribution to the control.

### S7 Zinc Finger nuclease treatment model

#### Visualisation of Zinc Finger concentrations during treatment

As explained in the ‘Methods’ section in the main text, we simulate the treatment of cells by mitochondrially targeted Zinc Finger Nucleases (mtZFNs). The concentration of the mtZFNs is modelled by the following equation:

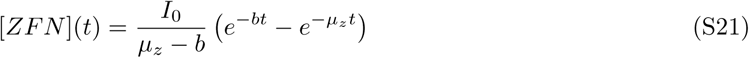

where *I_0_* is the treatment strength (e.g. the initially added mtZFN concentration), *b* indicates the treatment duration and *µ_z_* is the mtZFN degradation rate. Figure S6A shows this equation for different treatment durations. As we found in the main text, the weaker and longer treatment option leads to larger heteroplasmy shifts.

**Figure S6:**
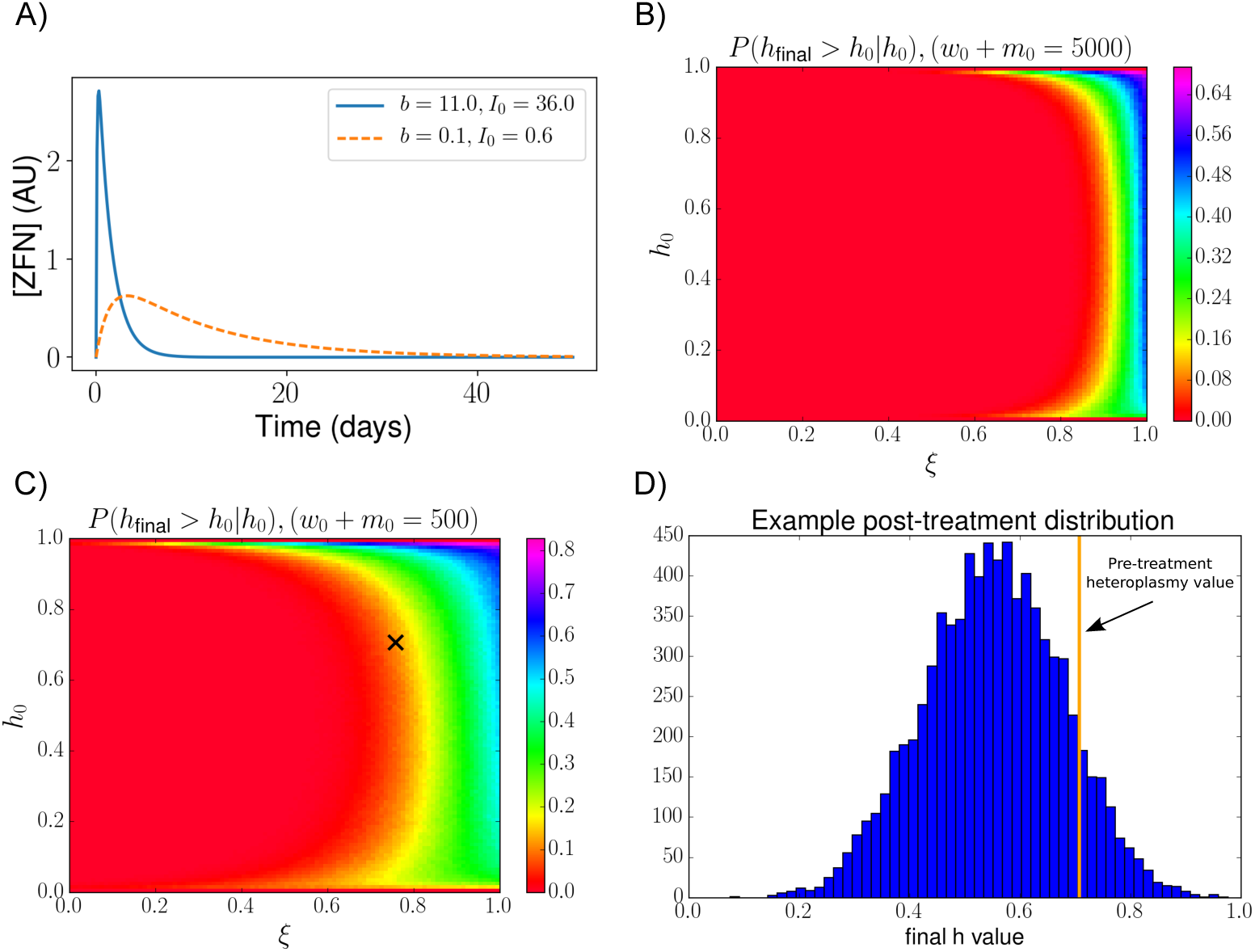
Zinc Finger Nuclease concentrations for short and long treatments. A) Here we show the concentration of mitochondrially targeted Zinc Fingers as modelled by equation (S21). The parameter values for the short and strong treatment (*I_0_* = 36,*b* = 11) are similar to those found in fitting the model to the data. For the mtZFN degradation rate we used *µ_z_* = log(2) (corresponding to a mtZFN half-life of 1 day). **There exists a possibility of increasing heteroplasmy levels through treatment. B)** The probability of increasing heteroplasmy above its initial pre-treatment value *h_0_*, after one round of treatment and recovery, is shown as a function of *h_0_* and ξ. Cells are initialized with a total copy number of 500. The cross indicates the parameters used in Figure (D). The parameter values for *I_0_, b* and *c_1_* are set to their fitted values: (*I_0_, b, c_1_) ≈* (39, 20, 3 × 10^−4^). Other parameters used are *µ_z_* = log(2) and δ = 1. C) Similar to figure (B), but now cells are initialized with a total copy number of 5000; in these large copy number cells stochastic fluctuations in copy number have less effect and the probabilities of exceeding initial heteroplasmy values are smaller compared to Figure (B). D) An example of a distribution of post-treatment heteroplasmy values is shown using parameters *h_0_* and ξ as indicated by the cross in Figure (B). The orange line indicates the value of *h_0_* (the heteroplasmy that was present before the treatment started).

#### Heteroplasmy values can increase after nuclease treatments

As mentioned in the Main text in Section 2.3.2, there is a possibility for cellular heteroplasmy to increase after a treatment has been applied. This is true especially if the selectivity of the treatment is low (i.e. ξ is close to 1) and the initial heteroplasmy of a cell is high; in this case treating a cell may even eliminate all wildtype mitochondria, increasing heteroplasmy to 1. To model the extent of this effect we initialize a cell with a given heteroplasmy *h_0_*, and let it undergo one round of treatment and recovery after which the final heteroplasmy is recorded. This process is repeated to obtain the probability that, given an initial heteroplasmy *h_0_*, the final heteroplasmy after treatment exceeds *h_0_* (P(h_final_ > *h*_0_|*h*_0_)). Figures S6B,C show these probabilities as a function of *h_0_* and selectivity parameter ξ, for initial mtDNA copy numbers 500 and 5000; the effect-size is larger for low copy number cells. Figure S6D shows an example of the distribution of post-treatment heteroplasmies. The recovery time used in the simulations is 30 days which is long enough for the cells to recover their initial copy numbers and short enough for the change in *h* to be almost completely due to treatment, rather than due to naturally occurring random drifts in heteroplasmy values. Chances of increasing heteroplasmy are highest when *h_0_* is very high or very low (if *h_0_* is low the low mutant copy numbers increase the effect of stochastic fluctuations). In the examples shown, when ξ <= 0.6 (i.e. for every mutant that is cleaved, 0.6 wildtypes are cleaved) increases in heteroplasmy are very unlikely to occur.

#### Details of Figure 5 in the Main text

Our fits to the experimental data [28] were obtained using deterministic (ODE) simulations and least square error. During the treatments a cellular linear feedback control *λ*(*w,m*) = c_0_ + c_1_(*w + δm*) is present in order for copy numbers to recover. Figures 5A and B in the Main text show deterministic and stochastic treatment trajectories using parameter values (*I_0_, b, ξ, c_1_) ≈* (40,12,0.76,5 x 10^−5^), which are the fitted parameters obtained when assuming the cells control the quantity *(w* + δ*m)* towards a value of 1000. Parameter c_0_ was set to be *c*_0_ = *µ* – *c_1_* (w_0_ + *δm_0_) = µ* – *c_1_* 1000 with *µ* = 0.07 corresponding to an mtDNA half-life of 10 days. Figure 5C assumes trajectories start in their deterministic steady state. The trajectories shown in Figure 5G have their treatment strengths optimized with respect to our cost function using the saturating model with low copy numbers.

#### MtZFNs were not able to shift high-heteroplasmy 143B cybrid cells

The possibility of shifting heteroplasmy levels by transfection with mtZFNs was shown in [28, 29]. To further test the capacity of mtZFN to shift heteroplasmy, mtZFN^2G^ (second generation) were transfected into human osteosarcoma 143B cybrid cells bearing 99% m.8993T>G mtDNA, a mutation associated with neuropathy, ataxia, and retinitis pigmentosa (NARP). All methods used are as described in [30]. All possible combinations of mtZFN monomer and control plasmid, including the non-mtDNA targeted PDE12 mtZFN, were transfected. Cells were subjected to FACS 24 hours post-transfection and cells expressing both constructs were sorted, total cellular DNA was extracted and mtDNA heteroplasmy and copy number measured (Figure S7A, B). Cell lines transfected with control plasmids or single mtZFN monomers did not demonstrate any shifts in heteroplasmy or changes in mtDNA copy number. Cells expressing the PDE12 constructs exhibited a depletion of copy number, to around 50% of control, but no shift in heteroplasmy was detected. Cells expressing dual mtZFN targeting the m.8993T>G mutation demonstrated a depletion of mtDNA copy number, to around 10% of control, with a modest shift in wild-type mtDNA heteroplasmy from 1% to 10% when quantified, though the signal-to-noise ratio of these quantifications is poor. However, this shift was not observed when cells were measured 14 days post-transfection, nor at 28 days, though mtDNA copy number had recovered to the control level (Figure S7C, D). These data indicate that mtZFN are not able to shift heteroplasmy in 143B cells bearing 99% m.8993T>G mtDNA. This is consistent with our predictions that it is difficult to decrease mutant mtDNA heteroplasmy in cells with very high initial mutant heteroplasmy. (Section 2.3.2).

**Figure S7:**
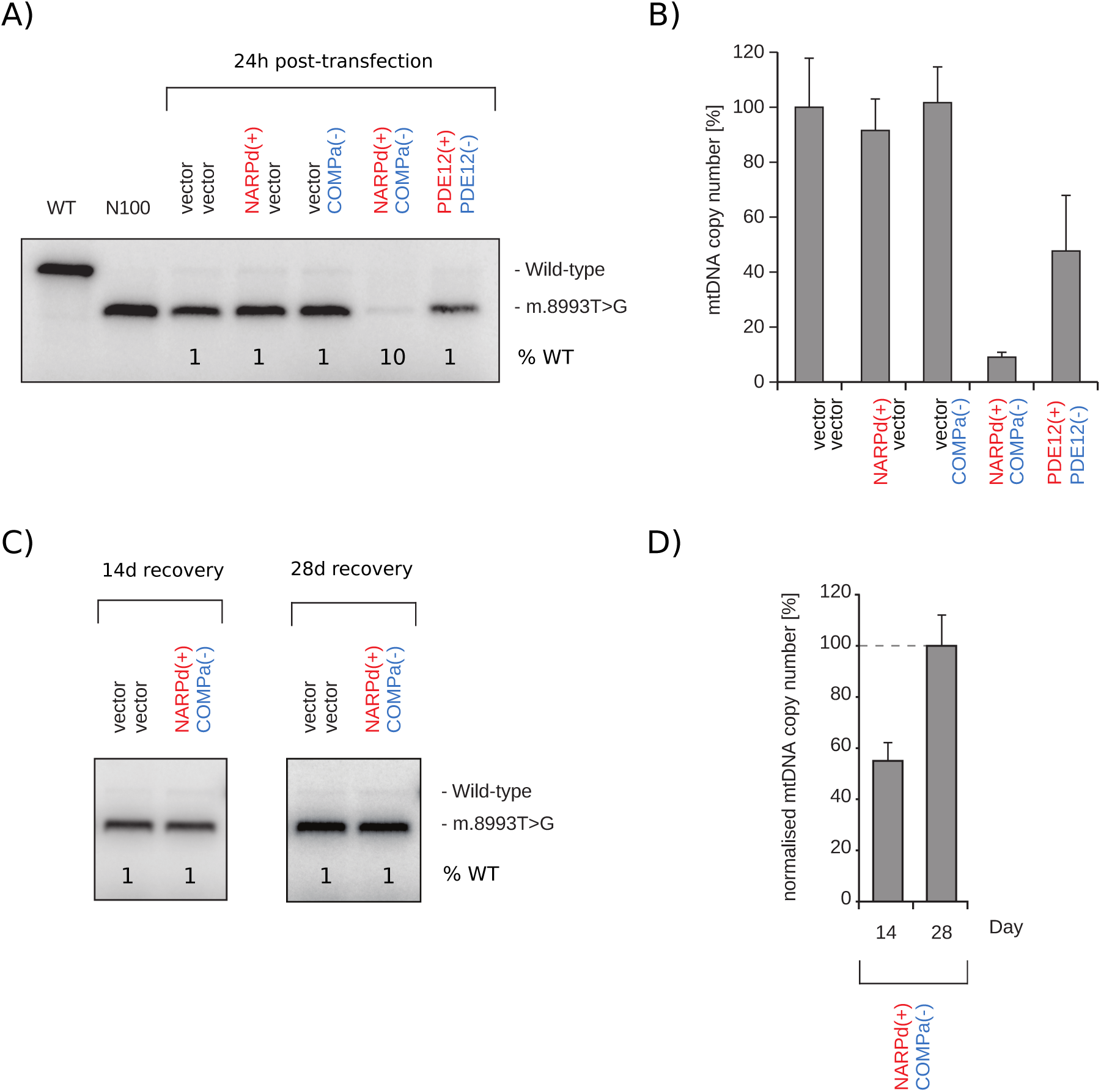
Effects of mtZFN^2G^ expression in a 143B cybrid cell line bearing 99% m.8993 T>G mtDNA. A), C) Last cycle hot PCR RFLP analysis of total cellular DNA extracts from cells transfected with indicated constructs and subjected to FACS sorting 24 hours (A), 14 days (C, left) or 28 days (C, right) post-transfection. PDE12 constructs are non-mtDNA sequence specific mtZFN used as a further control. B) qPCR measurement of mtDNA copy number in the same samples as (A). D) qPCR measurement of mtDNA copy number in the same samples as (C, left). For Figures (B) and (D) the experiments were performed in quadruplicate; error bars indicate 1 standard deviation.

#### Footnotes

1 Its effect on an arbitrary function *f(n)* is given by e.g. 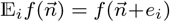 and 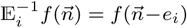 (where ϵ_1_ is a column vector with all zeros and a 1 at entry *i)*. Using a Taylor expansion, the operator can be written as 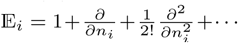.

2 𝔼_i_ changes n*i* into n*i* + 1, which is equivalent to changing ξ_i_ into ξ_i_ + Ω^−1/2^.

